# Glia mediate brain-wide activity and sleep behavior during sickness through an adrenergic-glutamatergic axis

**DOI:** 10.64898/2026.07.02.736189

**Authors:** Sara B Noya, Jianing Yang, Zhifeng Yue, Andrew Nguyen, Camilo Guevara, Shivani R Gianani, Vivian Xu, Pavel Pivarshev, Julie Williams, Kiet Lu, Amita Sehgal

## Abstract

Sickness induces coordinated changes in physiology and behavior, but despite the identification of circuits that underlie specific aspects of sickness, how the brain orchestrates this global state shift remains unclear. Whole-brain-single-cell RNA sequencing in a Drosophila model of sickness revealed that glia, rather than neurons, undergo the most extensive transcriptional remodeling and signal to neurons to drive sickness sleep behavior. Specifically, the process requires octopaminergic signaling and gap junctional coupling across a network of ensheathing glia (EG). EG modulate extracellular glutamate that is perceived by neurons via ionotropic glutamate receptors, producing a wave of neuronal activation that follows the EG calcium increase across the brain. Altogether, these findings elevate glia from passive supporters to active coordinators of sickness behavior, opening new avenues for understanding the neuromodulatory basis of sleep in the context of fatigue and malaise.

## Introduction

Sickness triggers a fundamental reorganization of physiology and behavior during which animals reduce their activity, redistribute energy, and enter a sleep-like state that promotes recovery (Davis and Raizen, 2017; Irwin, 2019; Kent et al., 1992). Studies have described humoral factors and sparse neuronal circuits that drive different components of sickness behavior (Darmohray et al., 2025; Ilanges et al., 2022; Jin et al., 2024; Konsman et al., 2002; Marvel et al., 2004; Osterhout et al., 2022a), however we are still missing how the brain coordinates a multimodal behavioral reconfiguration that is characterized by changes in activity across multiple brain regions.

Glia are placed at multiscale interfaces of the nervous system, wrapping synapses, shielding axons and neuropils and forming barriers with the periphery (Allen and Lyons, 2018; Fernandes et al., 2024; Freeman and Doherty, 2006; Kremer et al., 2017). In addition, glia form vast networks and their spatiotemporal span is found to be broader than ever expected (Chever et al., 2016; Oliveira et al., 2026; Orthmann-Murphy et al., 2007; Rash et al., 2001), which leaves them uniquely positioned to facilitate unified neural responses across the brain. Indeed, astrocytes participate in cortical synchronization and in the recruitment of neurons during hippocampal bursting and cortical UP states (Chever et al., 2016, 2016; Poskanzer and Yuste, 2016). Local neurotransmitter inputs result in broad astrocyte network activity changes (Cahill et al., 2024) and, even more, recent data indicate that the activation of these networks can persist across days as “astroengrams” (Dewa et al., 2025; Sánchez Romero and Navarrete, 2026). This raises the question of whether glia have instructive roles in state regulation under long-lasting conditions that involve brain-wide responses, such as sickness behavior. To date, the focus in such behaviors has been on neuron-centric mechanisms.

To determine how the brain coordinates a response in sickness, we profiled the adult Drosophila brain by single-cell RNA sequencing following a peripheral injury. Strikingly, glia showed the most profound transcriptional remodeling of any cell class, suggesting that glia are not passive responders but actively reprogrammed during sickness. Functional experiments reveal a glia-to-neuron axis in which adrenergic signaling through octopamine, the mammalian noradrenaline homolog, activates glia, and glia in turn modulate neurons through glutamatergic signaling to induce sleep. Interfering with adrenergic signaling in glia prevents glial and neuronal brain-wide activation and blocks sickness-associated sleep.

These results support a model in which glia integrate systemic sickness cues through adrenergic neuromodulation and transmit a brain-wide signal that reconfigures neuronal activity and promotes sickness sleep.

### Sickness sleep does not rely on canonical sleep-regulatory circuits

Flies exhibit changes in sleep amount during sickness, which can be triggered by challenges ranging from infection and neuronal damage to heat stress (Kuo et al., 2010; Lenz et al., 2015; Singh and Donlea, 2020). Even a peripheral injury, involving the injection of a micro volume of sterile PBS, produces whole-body transcriptional changes and an increase in sleep that dissipates after a maximum of 48h without affecting survival (Kuo et al., 2010; Troha et al., 2018) Thus, flies are an ideal model to understand the mechanisms underlying sickness sleep independently of the specifics of the disease etiology and progression. Since this model involves a breach in the fly cuticle, it is possible that ambient bacteria or the surface microbiome are recognized by the fly immune system. To eliminate this confound we monitored sleep in 7–10-day-old female *Canton-S* flies raised under sterile versus standard conditions and subjected to peripheral injury at ZT18 (see Materials and Method for rationale on the injury protocol). Regardless of the possibility of microbial recognition, flies exhibited indistinguishable responses to saline injections, with both axenic and standard groups displaying a significant increase in total sleep in the morning post injury (fig. S1A). As previously described (Kuo et al, 2010), this increase was observed during the 4h bin following lights on (fig. S1B) and was characterized by an increased bout number and bout length relative to sleep in uninjured flies at the same time mode (fig. S1C–D), indicating that sickness induced sleep is a consolidated rather than fragmented sleep.

We next asked whether sickness-induced sleep engages canonical sleep-regulatory pathways so we evaluated the sickness response of low-sleep mutants—*sleepless* (*sss^p1^*, *sss^p2^*) and *fumin* (*fmn*)— that carry neuronal defects underlying their sleep phenotypes (Koh et al., 2008; Kume et al., 2005). Following peripheral injury, *sss^p1^*and *fmn* mutants showed an immediate increase in sleep after injury (fig. S1E and F top, blue and orange). *sss^p2^* flies, which maintain normal baseline sleep, but have homeostatic sleep defects, exhibited the canonical morning sickness sleep (fig. S1E and F bottom, dark red). This indicates that sleep induced by a peripheral injury does not rely on these classic sleep mechanisms. We then tested whether other sleep-promoting pathways contribute to sickness sleep. Acute optogenetic inhibition of canonical sleep-promoting neurons (23E10-GAL4 driven circuits) (De et al., 2023; Jones et al., 2025) or perturbation of lipid-transfer between glia and neurons (*repo*-GAL4>UAS-*GLaz* RNAi), which has recently emerged as a key element in sleep modulation (Haynes et al., 2024; Pyfrom et al., 2025), also failed to reduce injury induced sleep (fig. S1G and H).

Collectively, these data support a model in which sickness sleep is distinct from classic homeostatic sleep and suggest the involvement of a dedicated neuronal mechanism to promote sleep during physiological stress.

### Single-cell profiling reveals the largest transcriptional changes in glia after peripheral injury

Growing interest in sickness behavior has led to the mapping of brainstem and hypothalamic circuits that are activated following a peripheral immune challenge (Darmohray et al., 2025; Ilanges et al., 2022; Jin et al., 2024; Marvel et al., 2004; Osterhout et al., 2022b). These observations occur in the context of widespread changes across the brain (Ilanges et al., 2022; Marvel et al., 2004), but how the brain coordinates its response across diverse cell types and nuclei remains elusive. To address this gap, we leveraged the simple cellular architecture of the *Drosophila* brain and performed single-cell RNA sequencing to obtain an unbiased view of cell types and gene programs altered during sickness (an interactive online resource was generated to query gene expression changes across cell types (bit.ly/3PaWENn).

We collected brains after a sterile injury at the beginning of the sickness sleep episode (Fig. 1A) and recovered 7,914 cells (53% control, 47% injured) and ∼8,000 detected genes with sequencing depth comparable to similar datasets (Davie et al., 2018; Dopp et al., 2024) (52k reads per cell; median 1.8k genes). Dimensionality reduction (t-SNE, UMAP) clearly separated neurons (∼80%, *nSyb*⁺) from glia (∼20%, *repo*⁺) (fig. S1I). Iterative Leiden clustering and multi-step annotation assigned 3,094 cells (39%) to 13 clusters spanning four glial types and several neuronal classes (fig. S1J and K). To overcome the low power of clusters with small cell numbers (Liu et al., 2023), we grouped neurons by the expression of the main neurotransmitter. This resulted in four major neuronal types—cholinergic, glutamatergic, gabaergic, and monoaminergic (Fig. 1B and C). Glia were divided into two large clusters, astrocyte-like cells and ensheathing glia, and two small clusters, cortex glia and a group of unclassified glia cells (Fig. 1B and C, fig. S1J and K). Both neurons and glia contained similar proportions of control and injured cells (Fig. 1D).

**Figure 1.**
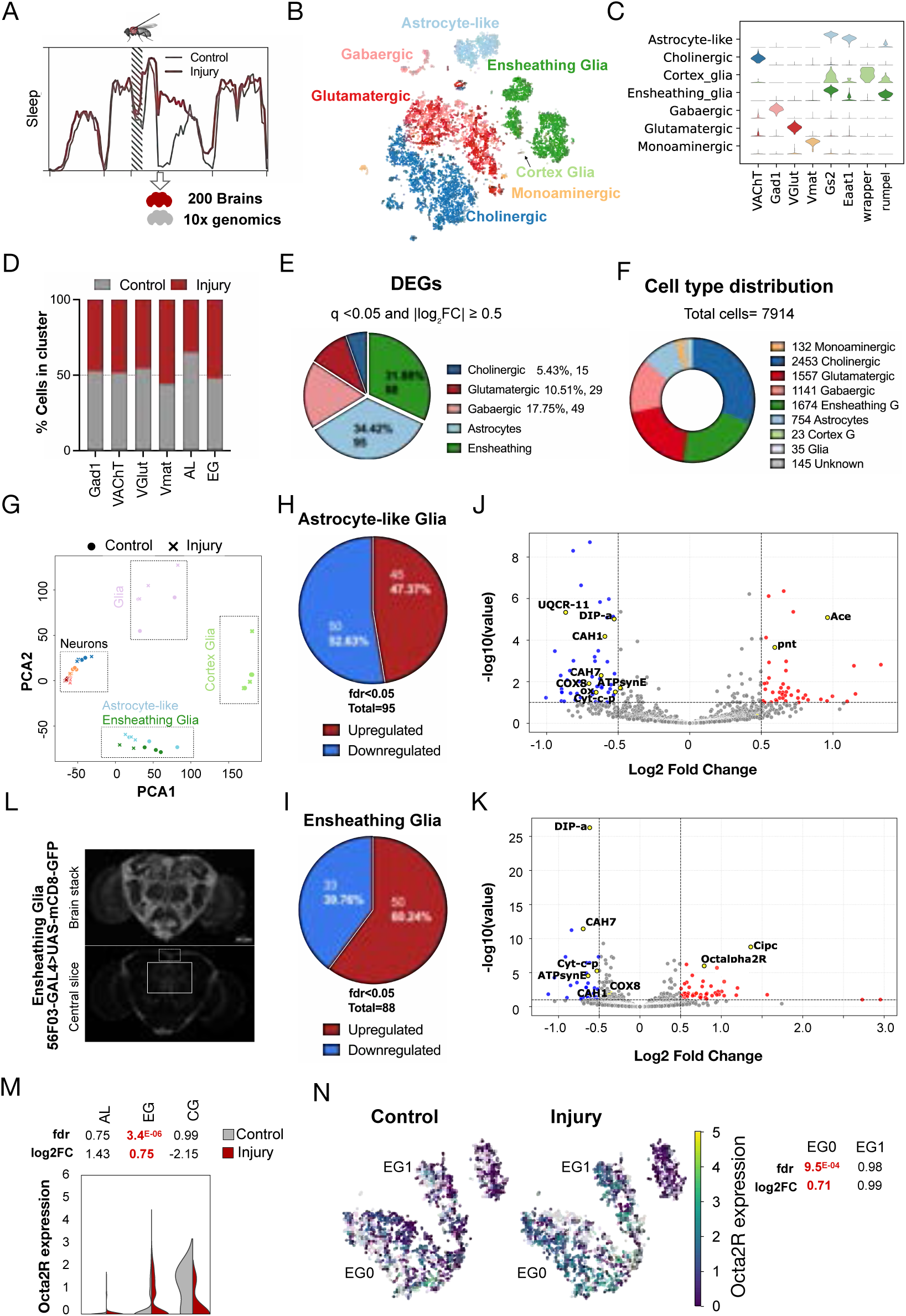
Glia undergo the largest transcriptional changes after a peripheral injury. (A) Schematic for scRNAseq protocol of Drosophila brains during sickness. Flies were injured in the middle of the night and 6h later, when control and experimental groups display the maximal difference in sleep, fly brains were dissected, dissociated and sorted to remove dead cells and debris. Library preparation was performed on the same day, followed by 10x Genomics sequencing. A custom analysis pipeline base on Scanpy guidelines was applied for data analysis (See Materials and methods for details). (B and C) tSNE (B) and marker gene expression (C) of 7k cells of 5-day-old fly brains (controls and injury), colored by main expression of *VAChT* indicating cholinergic neurons in dark blue, *VGlut* indicating glutamatergic neurons in red, *Gad1* indicating GABAergic neurons in pink, *Vmat* indicating peptidergic neurons in orange. Astrocytes in light blue were classified by expression of *Eaat1,* ensheathing glia in dark green by the expression of *rumpel* and cortex glia in light green by the expression of *wrapper*. (D) Proportion of cells from injured and control flies from clusters represented in (B and C). (E) Pie chart indicating the number of DEGs per cluster. DEGs were selected based on a fdr < 0.05 and log fold change lower than −0.5 or greater than 0.5. (F) Proportion of cells per cluster. (G) PCA of pseudobulb samples from control (circles) or injury (x) flies, color coded by cell type. (H-K) Pie charts and volcano plots (J and K) of upregulated (red) and downregulated (blue) genes in Astrocyte-like glia (H and J) and ensheathing glia (I and K). DEGs are highlighted in blue (downregulated) and red (upregulated) and include genes with fdr < 0.05 and log fold change lower than−0.5 or greater than 0.5. Highlights in yellow correspond to genes mentioned in the text. (L) Immunofluorescence staining of ensheathing glia using membrane bound GFP (56F03-GAL4>UAS-mCD8-GFP). Top is the entire brain projection and bottom is a slice capturing the ensheathment of the central complex. Note the ensheathment of cell bodies (dashed square) and the neuropil (straight square). (M) Violin plot showing expression of Oct2aR in control (grey) and injured (red) samples in the glia clusters. Expression data correspond to log1p normalization. (N) tSNE of EG subclusters indicating expression of Oct2aR in control (left) and injury (right) cells. Expression data correspond to log1p normalization. Marker gene expression for the subclusters can be found in Figure S2A and B.

Differential expression analysis revealed modest number of significantly changed genes in most neuronal classes. Using q <0.05 and |log₂FC| ≥ 0.5, cholinergic neurons showed 15 differentially expressed genes (DEGs), glutamatergic cells showed 29 and gabaergic had 49 (Fig. 1E and Table S1). Interestingly, a few genes changed consistently across neuronal classes: the axon-guidance receptor *robo2* and the metabolic enzyme pyruvate carboxylase (*Pcb*) were broadly downregulated, while *Hsp23* was consistently upregulated (fig. S1L-N), in agreement with metabolic reorganization and a cellular stress response.

Glia, despite representing only one-third of the total cells, accounted for roughly two-thirds of all significant DEGs (Fig. 1E and F). We wondered if this could be due to cluster homogeneity so we measured cluster entropy which accounts for purity and trajectory analysis. Despite glia having slightly lower entropy, when considering cluster size, we found that all clusters were close to Shanon maximal entropy, so this does not appear to explain the differences in DEG numbers. PCA further revealed a marked separation of control and injured pseudobulk replicates from astrocytes, ensheathing glia, and the unclassified glial cluster (Fig. 1G). By contrast PCA distances were smaller in neuronal samples, consistent with a pronounced and coordinated glial response.

Astrocytes and ensheathing glia exhibited widespread downregulation of genes encoding components of the mitochondrial electron transport chain—including *SdhC*, multiple cytochrome c subunits (*Cyt-c-p*, *COX7a*, *COX8*), complex III (*ox, UQCR-11*), and several complex I genes—suggesting reduced oxidative phosphorylation and increased ROS burden (Fig. 1H-K). Also diminished were the carbonic anhydrases *CAH1* and *CAH7* (Fig. 1H-K), that are modulated in response to CO₂ and pH changes in neuroprotective contexts (Theparambil et al., 2024). By contrast the Acetylcholine esterase *Ace*, the senescence transcriptional factor *pointed* (*pnt*) and the clock interacting protein circadian (*cipc*) increased in astrocytes and ensheathing glia respectively (Fig. 1H-K).

Taken together, these findings reveal that glial cells—not neurons—undergo the strongest transcriptional shifts after peripheral injury, suggesting that glia may play a key and previously underestimated role during sickness behavior.

### Noradrenergic signaling promotes sickness-induced sleep through *Octα2R* in ensheathing glia

Ensheathing glia (EG) and astrocytes populate the neuropil, where they support metabolism and maintain ionic and neurotransmitter balance (Bittern et al., 2021; Kremer et al., 2017; Yildirim et al., 2019). Both cell types have been linked to sleep regulation (Blum et al., 2021; Flores-Valle et al., 2025; Stahl et al., 2018; Vanderheyden et al., 2018). Ensheathing glia specifically form continuous networks that surround brain compartments (Fig. 1L); therefore, they are perfectly positioned to propagate and coordinate changes across the brain. To identify pathways that enable glia to respond to injury, we searched for surface receptors upregulated in EG and astrocytes in our scRNAseq dataset. The octopamine receptor *Octα2R*, a homolog of mammalian adrenergic receptors that has been recognized for its effect on locomotion (Nakagawa et al., 2022), was one of the top upregulated genes in EG (Fig. 1K and Table S1). Although expressed across cell types, significant upregulation of *Octα2R* was unique to EG (Fig. 1M and Table S1). Even more, the differential expression was specific for a distinct EG subclusters characterized by the expression of the transient receptor potential channel *waterwitch* (*wtrw)* (EG0 subcluster) (Fig. 1N, fig. S2A and B and Table S2).

We next asked whether *Octα2R* contributes to the sleep increase that follows a peripheral injury. Knocking down *Octα2R* in EG (56F03-GAL4,UAS-*Octα2R* RNAi) abolished the characteristic post-injury sleep enhancement observed in controls (Fig. 2 A and B and fig. S2C. See fig. S6 for KD efficiency). We also knocked down *Octα2R* in astrocytes (*alrm*-Gal4) and cortex glia (NP2222-Gal4) but none of those manipulations reduced the injury-evoked response (Fig 2C and D and fig. S2D). Interestingly, KD in surface glia (9-137-Gal4) didn’t result in significant differences in sleep during the 4h window we use to evaluate sickness sleep but in a 2h delayed window (ZT0-6) showed a clear trend to increased sleep time in injured versus control flies behavior (Fig. 2C and fig. S2E).

**Figure 2.**
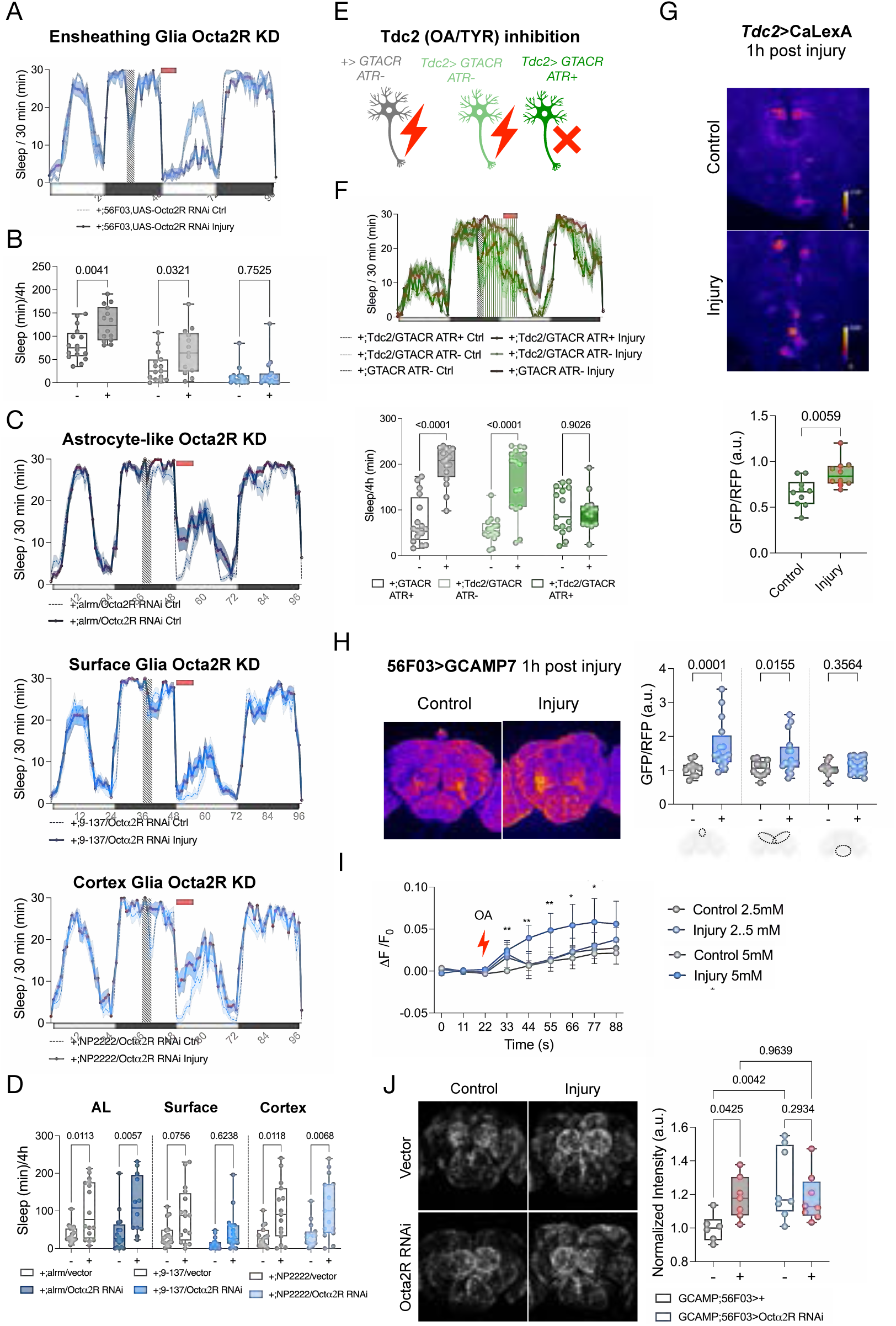
Ensheathing glia activation through *Octα2R* is required for sickness sleep. (A and B) Sleep profile after peripheral injection of flies with adult-specific knockdown of Oct2aR in ensheathing glia (blue). See fig. S2C for genetic controls. For all sleep experiments, the sleep profiles correspond to the average time spent sleeping in consecutive 30-min segments; mean. Thin dashed lines correspond to handled control flies and thick lines with red symbols to injured. Time of injury is indicated by stripped bar and sickness sleep window (ZT0-4) by a red horizontal bar. (B) Quantification of time spent asleep between ZT0–4 for flies from A. Two-way ANOVA was followed by Tukey’s correction. n=10-16. For clarity, only the control and injury comparison for each genotype are shown. See Table S2 for full statistics. (C and D) Sleep profile after peripheral injection of flies with adult-specific knockdown of Oct2aR in astrocytes (Alrm-GAL4), cortex glia (NP2222-GAL4) and surface glia (9-173-GAL4). See fig. S2D for genetic controls. (B) Quantification of time spent asleep between ZT0–4 for flies from A. For the surface glia KD (9-13-GAL4), an additional quantification is show in fig.S2E corresponding to the ZT0-ZT6 time window. ANOVA was done, applying a Two-way ANOVA followed by Sidak multiple testing correction. Comparisons of the control and injured condition for each genotype are shown. n=10-16. (E) Diagram of the experimental design for the Tdc2 optogenetic activation. In flies expressing both the GAL4 and the UAS, lack of ATR prevents the optogenetic activation of neurons producing TYR and OA. In flies fed on ATR (ATR+) when exposed to light, Tdc2+ neurons get activated. (F) Sleep profile after peripheral injection of flies in which Tcd2 neurons were optogenetically silenced (dark green) and respective genetic controls (grey corresponds to the genetic control pre fed with ATR and light green correspond to the experimental genotype fed with a vehicle). Vertical lines indicate the optogenetic activation (10s on 10 s off, See Materials and methods for details). (D) Quantification of time spent asleep between ZT0–4 for flies from C. Two-way ANOVAs followed by Tukey’s correction. n=10-16. (G) Calcium signal in Tdc2 ventral cells expressing CaLexA (fire LUT) (See S2F for entire brain image and RFP internal control). (Bottom) Quantification of Tdc2+ cells. Unpaired two-tailed t-test. n=10. (H) Ca imaging and qualification of ensheathing glia 1h post injury, while driving the expression of CaLexa with the 56F03-GAL4 driver. For analysis, brain was subdivided into a PI region, central brain and ventral brain (see diagram). See S2G for internal control expressing RFP. Data are normalized to the average of the control brain data for each region. ANOVA with Holm-Sidak multiple correction. n=10-16. (I) Ca levels in EG after bath application of 2.5 and 5 mM of octopamine (OA). Brains were dissected 1h after injury and imaged in a saline bath containing 1uM TTX to block neuronal activity. ANOVA with Sidak multiple correction. Stars correspond to the control-Injury comparison in the 5mM condition. n=4. *<0.01,**<0.001. See Table S2 for full statistics. (J) Ca signal in ensheathing glia 1h post injury, while driving the expression of the genetically encoded Ca indicator GCAMP7 with the 56F03-GAL4 driver in flies with Oct2aR knockdown (blue), compared to the genetic controls expressing an empty vector (grey). Data are normalized to the average of the control. Two-way ANOVAs followed Tukey’s correction. n=5-8.

The previous finding suggests a crucial role of octopaminergic signaling acting in EG to modulate sickness sleep. In the fly, octopamine synthesis depends on a two-step process that involves the Tyrosine decarboxylase 2 (Tdc2) and Tyramine β-hydroxylase (*Tbh*) enzymes (Fig. 2E top) so we tested whether *Tdc2*⁺ neurons are necessary for injury-induced sleep. Silencing *Tdc2*⁺ neurons using the light-gated chloride channel GTACR (Fig. 2E top) suppressed the sleep increase normally seen after injury (Fig. 2E and F), corroborating that adrenergic signaling is required during sickness behavior. Indeed calcium imaging of *Tdc2*⁺ neurons using the calcium indicator Calexa showed that a ventral subset, shown to co-express *Tbh* (Schneider et al., 2012), had increased calcium, indicative of higher activity, in injured flies compared to controls (Fig. 2G and fig. S2F).

Thus, ensheathing glia emerge as a critical player in the manifestation of sickness behavior, mediating a noradrenergic axis that is activated after injury.

### Injury activates EG across the brain through *Octα2R*

*Tdc2* neurons release both tyramine and octopamine to drive astrocytic calcium increases (Ma et al., 2016). Because astroglial calcium dynamics appear to encode homeostatic sleep need in both flies and mammals (Blum et al., 2021; Flores-Valle et al., 2025; Ingiosi et al., 2020), we hypothesized that octopamine-driven calcium in EG may similarly encode sickness-induced sleep pressure. Using the intracellular Ca indicator CaLexA expressed in EG (56F03>CaLexA), followed by brain regional analysis, we observed increased Ca in the dorsomedial and central brain 1 h post-injury, but detected no differences at 6 h (Fig. 2H and I; fig. S2G and H). Similarly, imaging of brain *ex vivo* (56F03>GCaMP7) revealed an increase in EG calcium 1h post-injury (fig. S2I). At 6h post injury, coinciding with the morning peak of locomotor activity, both control and injured flies increased calcium significantly and differences were no longer noticeable between groups (fig. S2I); similar increases in calcium at this time were previously shown in neurons (Cuddapah et al., 2025). Thus, normal wake and sickness sleep both correlate with increased calcium activity in EG across the brain, which highlights the need to better characterize the mechanistic differences and implications of calcium dynamics across different behavioral states.

We then tested the hypothesis that OA activates EG. In ex vivo preparations in a bath containing tetrodoxin (TTX) to eliminate neuronal OA output, octopamine (OA) application triggered EG calcium increases at both 2.5 and 5mM concentrations in control and injured brains (Fig. 2I). Interestingly, at the higher concentration, brains from injured flies exhibited heightened sensitivity with a steady and significant increase compared to any other group (Fig. 2I). By contrast, tyramine (TYR) did not elicit the same responses (fig. S2J). Along these lines, knockdown of a Tyramine receptor (*Oct-TyrR*), that was previously shown to participate in behavioral modulation through astrocytes (Ma et al., 2016), did not influence sickness sleep (fig. S2K), consistent with a specific role for OA and *Octα2R* rather than TYR in modulating sickness sleep.

Finally, to test if EG activation after injury is indeed dependent on octopaminergic signaling through *Octα2R*, we simultaneously knocked down the receptor and measured intracellular Ca. Interestingly, knockdown of Octα2R increased baseline levels of calcium in EG and there were no further increases induced by injury (Fig. 2J). In summary, this body of experiments indicates that OA modulates EG activation through Octα2R and that the increased expression of Octα2R during injury primes EG to respond to OA neuromodulation.

### Noradrenergic signaling modulates glutamate homeostasis through EG

Recent work has highlighted how catecholamines can engage astrocytes to influence neuronal output through modulation of extracellular adenosine (Chen et al., 2025; Guttenplan et al., 2025; Lefton et al., 2025). EG, on the other hand, participate in glutamate homeostasis in the brain, through which they encode negative valence during memory acquisition (Miyashita et al., 2023; Otto et al., 2018). To test if glutamate dynamics change in injured brains, we expressed the extracellular glutamate sensor iGluSnFR in EG (56F03>iGluSnFR). Extracellular glutamate increased 1 h after injury (Fig. 3A). However, 6 h post injury, glutamate interestingly diminished in injured flies (Fig. 3A).

**Figure 3.**
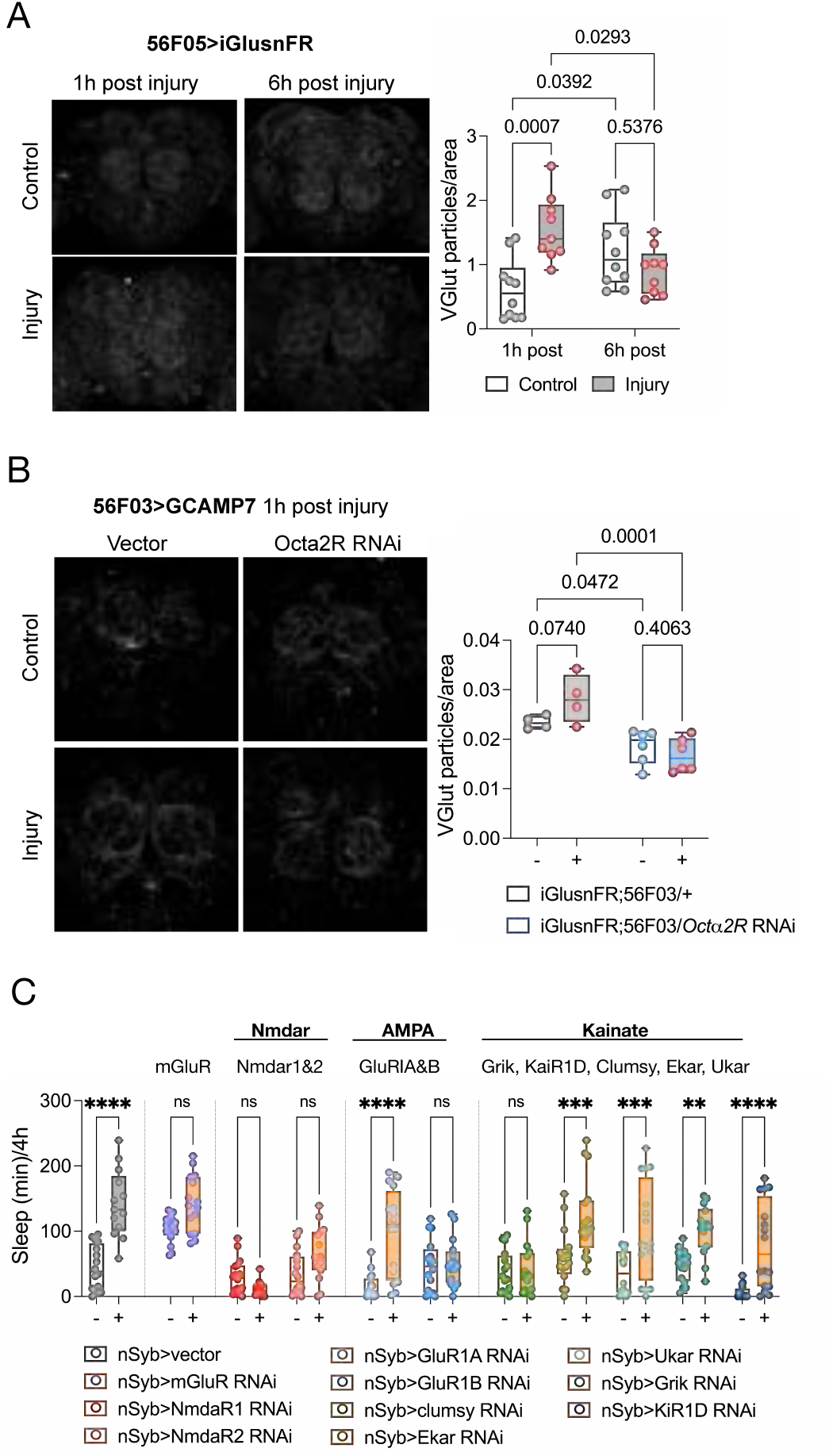
Noradrenergic signaling in EG modulates extracellular glutamate. (A) Images and quantification of fly brains expressing iGlusnFR under the control of an EG driver (56F03-GAL4). Images were taken at 1h (A left) and 6h (A right) post injury. Glutamate signal was quantified and normalized to the handled control flies at 1h post injury. To-way ANOVA followed by Sidak multiple testing correction. (B) Images and quantification of fly brains expressing iGlusnFR under the control of an EG driver (56F03-GAL4) with (blue) or without (grey) Oct2aR KD. Images were taken at 1h post injury. Two-way ANOVA followed by Sidak multiple testing correction. (C) Quantification of time spent asleep between ZT0–4 of knockdown of glutamate receptors in neurons (nSyb-GAL4). See fig. S3 for sleep profiles. ANOVA was done specifying the comparisons between control and injury for each genotype and Sidak multiple testing correction was applied. n=16,**<0.01, ***<0.001, ****<0.0001.

To directly test if this glutamate modulation was dependent on OA signaling through *Octα2R* we knocked down the receptor in EG while simultaneously expressing iGluSnFR. Measurement 1h post injury showed that KD of the receptor prevented an increase in extracellular glutamate (Figure 3B). In addition, EG knockdown of several genes involved in glutamate transport (*Eaat1* and Vglut1) and metabolism (*Gdh* and *Got2*) increased baseline sleep and reduced sickness sleep such that it was difficult to distinguish control and injured flies (fig. S3A). This aligns with the idea that EG dependent glutamate modulation is key to regulating morning sleep and perhaps sickness-induced sleep. Indeed, pan-neuronal KD of the metabotropic receptor *mGluR* KD increased baseline sleep to ceiling levels during the effective sickness window (Fig. 3C and fig.S3B).

In addition to *mGluR,* we knocked down other glutamate receptors in neurons to identify those that mediate the response to glutamate from EG. Of all glutamate receptors in the Drosophila genome, KD of the ionotropic receptors Nmdar1but, Nmdar2 (Fig. 3C and fig. S3B, see fig. S6A for KD efficiency) and Clumsy (GluR1B showed variability between replicates) blocked sickness sleep behavior without affecting baseline sleep indicating that glutamate is at least one of the mechanisms of the glia-neuron crosstalk during sickness.

These imaging and behavioral findings support a model in which ensheathing glia contribute to glutamate homeostasis and regulate extracellular glutamate downstream of noradrenergic signaling to modulate both morning and sickness sleep. The dissociable receptor requirements underlying these effects further suggest that glutamatergic signaling engages distinct and dedicated pathways to control specific sleep states.

### EG modulates neuronal activation during sickness to induce sleep

Recent mammalian studies have reported activation of most of the brain after a peripheral immune challenge (Ilanges et al., 2022; Marvel et al., 2004). To test if the increase in glial Ca in Drosophila during sickness behavior is also accompanied by brain-wide neuronal activation, we performed calcium dependent activity mapping using CaLexA under the control of a pan neuronal driver (*nSyb*-GAL4). 1h post injury, calcium was slightly higher in multiple brain neuropils (fig S4A). By 6h the differences between control and injured flies increased even further (Fig. 4A). This increased Ca in neurons was also observed in live flies using 2 photon microscopy (fig. S4B). By 12h, control and injured brains were indistinguishable (fig S4C). To test if the rise in Ca was causal, rather than collateral, we conditionally overexpressed in neurons the mammalian protein parvalbumin (He et al., 2018). This manipulation prevents calcium increases in ex vivo preparations of adult fly brains when expressed conditionally for 48h under the neuronal promoter *nSyb* (fig. S4D). When we conditionally expressed PV in neurons for 48h pre injury, sickness behavior was abolished (Fig. 4B). And although PV expression reduced daytime and nighttime sleep when expressed in three well established sleep promoting neuronal subsets (fig. 4SE), its expression in these neurons was not enough to abrogate sickness sleep (fig. S4F), indicating that these sleep centers are dispensable for sickness sleep behavior and supporting the idea that sickness sleep is different from daily or homeostatic sleep.

**Figure 4.**
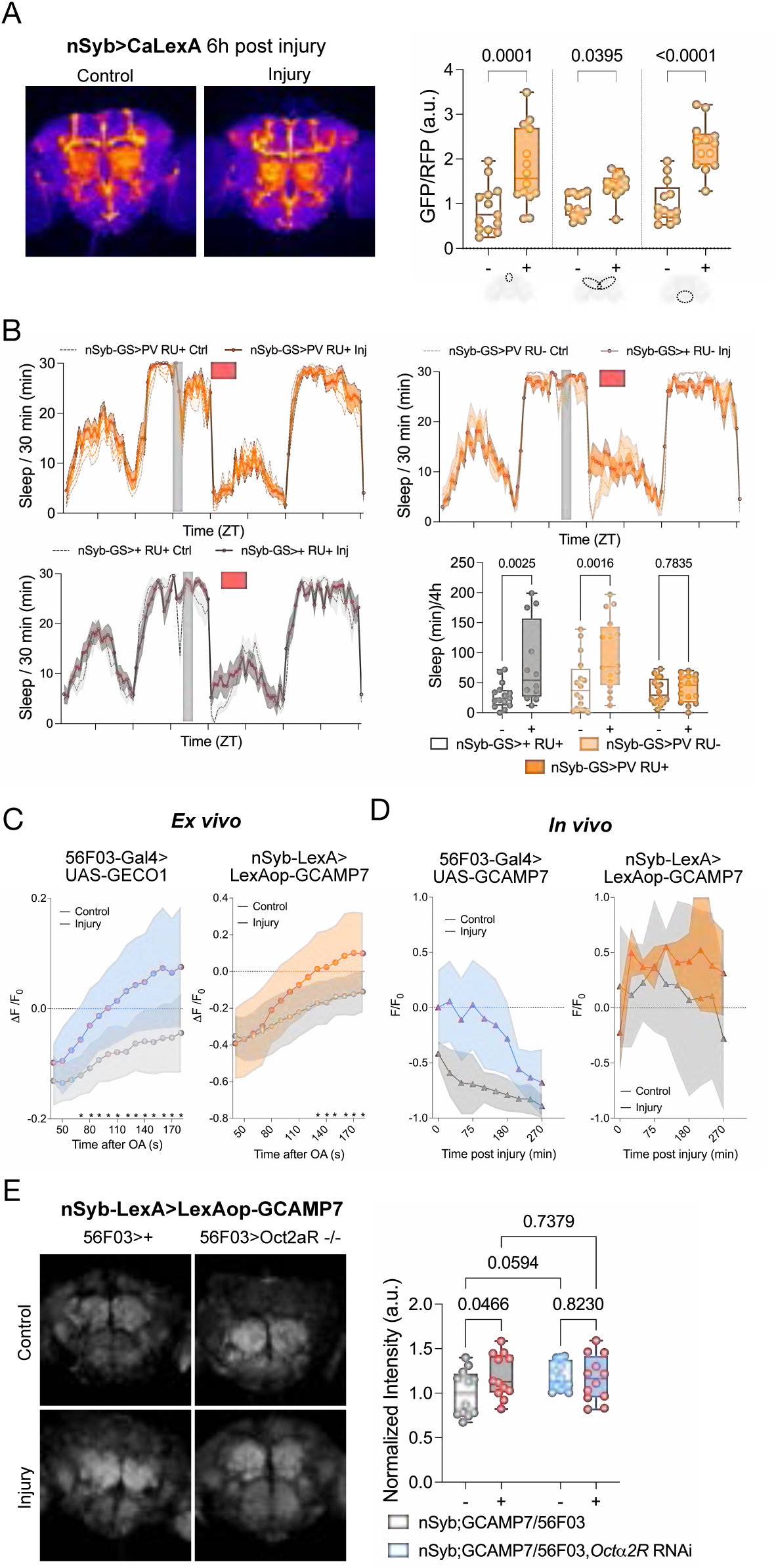
Brain-wide activity of EG and neurons during sickness sleep. (A) Ca imaging and quantification in neurons 1h post injury, while driving the expression of CaLexA with the nSyb-GAL4 driver. For analysis, brain was subdivided into a PI region, central brain and ventral brain (see diagram). ANOVA with Holm-Sidak multiple testing correction. n=10-16. (B) Sleep profile after peripheral injection of flies with adult-specific expression of parvalbumin (PV) in neurons (orange) and genetic controls (grey). Bottom right shows the quantification of time spent asleep between ZT0–4. ANOVA was done applying a Tukey multiple testing correction. For clarity, only the control and injury comparisons for each genotype are shown. No significant difference was found between sleep of handled control flies from the different genotypes. n=16. (C) *Ex vivo* measurement of Ca changes in EG (56F03-LexA>LexAop-GECO1, left) and in neurons (nSyb-GAL4>UAS-GCAMP7, right) in control (gray) and injured (color) brains. Measurements of neuronal and EG Ca were done with simultaneous acquisition after bath application of 5 mM of octopamine. Brains were dissected 1h after injury. ANOVA with Sidak multiple testing correction. n=8. (D) *In vivo* 2P imaging of Ca using GCAMP7 in ensheathing glia (lefts) and neurons (right). Optical windows were opened at least 2 days prior to the experiment. Flies were mounted and imaged every 30 min for 6 h after sterile injury in the thorax. n=3. (E) Ca signal in neurons 1h post injury, while driving the expression of the genetically encoded Ca indicator GCAMP7 with the nSyb-LexA (nSyb-LexA>LexAop-GCAMP7) driver in flies with simultaneous KD of Oct2aR in ensheathing glia (56F03-GAL4>UAS-Oct2aR RNAi) (blue), compared to the genetic controls expressing an empty vector (grey). Data are normalized to the average of the control +/+. ANOVAs followed Tukey’s multiple testing correction. n=10-12.

Since EG activation was higher at 1h post injury compared to 6h (Figure 2H-I and fig. S2G) whilst the largest neuronal Ca changes were observed 6h post injury (Figure 4A and fig. S4A), we wondered whether *Octα2R* signaling through EG is required for the peak in pan neuronal activation. *Ex vivo* simultaneous imaging of Ca signals in EG and neurons, revealed octopamine-evoked calcium increases in both neurons and glia (Figure 4C and fig. S4G). Increases in EG were measured within the first 10s of octopamine application while it took over 2 minutes to see significant increases in neurons. Although these differences could be due to the intrinsic properties of the indicators, our observation agrees with a previous study reporting that lower concentrations of OA trigger Ca increases in both astrocytes and EG as compared to neurons despite inducing smaller fold changes (Černe et al., 2025). In addition, it aligns with the idea that EG activation precedes panneuronal firing. We also monitored EG and neuronal calcium in live flies after injury. To avoid the confound of tissue damage that occurs during optical window preparations, we performed microsurgeries that allow imaging of neuronal activity for up to several days (Huang et al., 2018; Zhu et al., 2026). Two days after surgery, when sleep is normalized (Huang et al., 2018), we selected flies that had no noticeable defects in movement, set them into an imaging holder for 2 photon imaging and recorded activity over a span of 6h. We observed no differences in intracellular Ca in neurons between control and injured flies during this time window (Fig. 4D). By contrast, in EG, there was a net increase in injured versus control flies (Fig. 4D).

Lastly, to test if neuronal activation depends upon adrenergic signaling through EGs, we simultaneously knocked down *Octα2R* in EG while measuring neuronal Ca. As we observed for intracellular EG Ca (Fig. 2K), KD of *Octα2R* elevated baseline Ca levels in neurons and prevented further increases after injury (Fig. 4E).

Collectively, these findings delineate a glia-to-neuron signaling axis in which octopamine-activated EG modulates neuronal activation to coordinate the sleep response to injury.

### Gap junctional communication between EG is required for neuronal activation and sickness sleep

Given the temporal and spatial responses uncovered in the previous experiments we wondered if EG were propagating a signal across the brain similar to the astrocytic gap-junction-coupled networks that contribute to long-range functional connectivity (Charles, 1998; Cornell-Bell et al., 1990; Handy and Borisyuk, 2023; Peng et al., 2023). In Drosophila, gap junctions are formed by innexins. Inx2 is expressed in both astrocytes and ensheathing glia whilst Inx3 is primarily expressed in ensheathing glia (Davie et al., 2018; Li et al., 2022). Interestingly, Inx3 and Inx2 expression increases during sleep in EG (fig. S5A, adapted from https://www.flysleeplab.com/scsleepbrain<u>, (Dopp et al., 2024)</u>). Consistent with the idea that an EG network is necessary for sickness sleep behavior, we observed that knockdown of the EG-specific gap junction protein Inx3 impaired sickness sleep (Fig. 5A, See fig. S6 for KD efficiency). Inx2 KD resulted in high baseline sleep, but there was still a trend towards further increase (fig. S5B). To determine if the EG network also modulates the whole brain activation, we knocked down Inx3 in EG while simultaneously evaluating calcium in neurons (nSyb-LexA; LexAop-GCAMP7, 56Fo3-GAL4, UAS-Inx3 RNAi). As expected, flies with intact Inx3 showed increased neuronal Ca 1h post injury but Inx3 KD prevented the neuronal calcium activation after injury (Fig. 5B).

**Figure 5.**
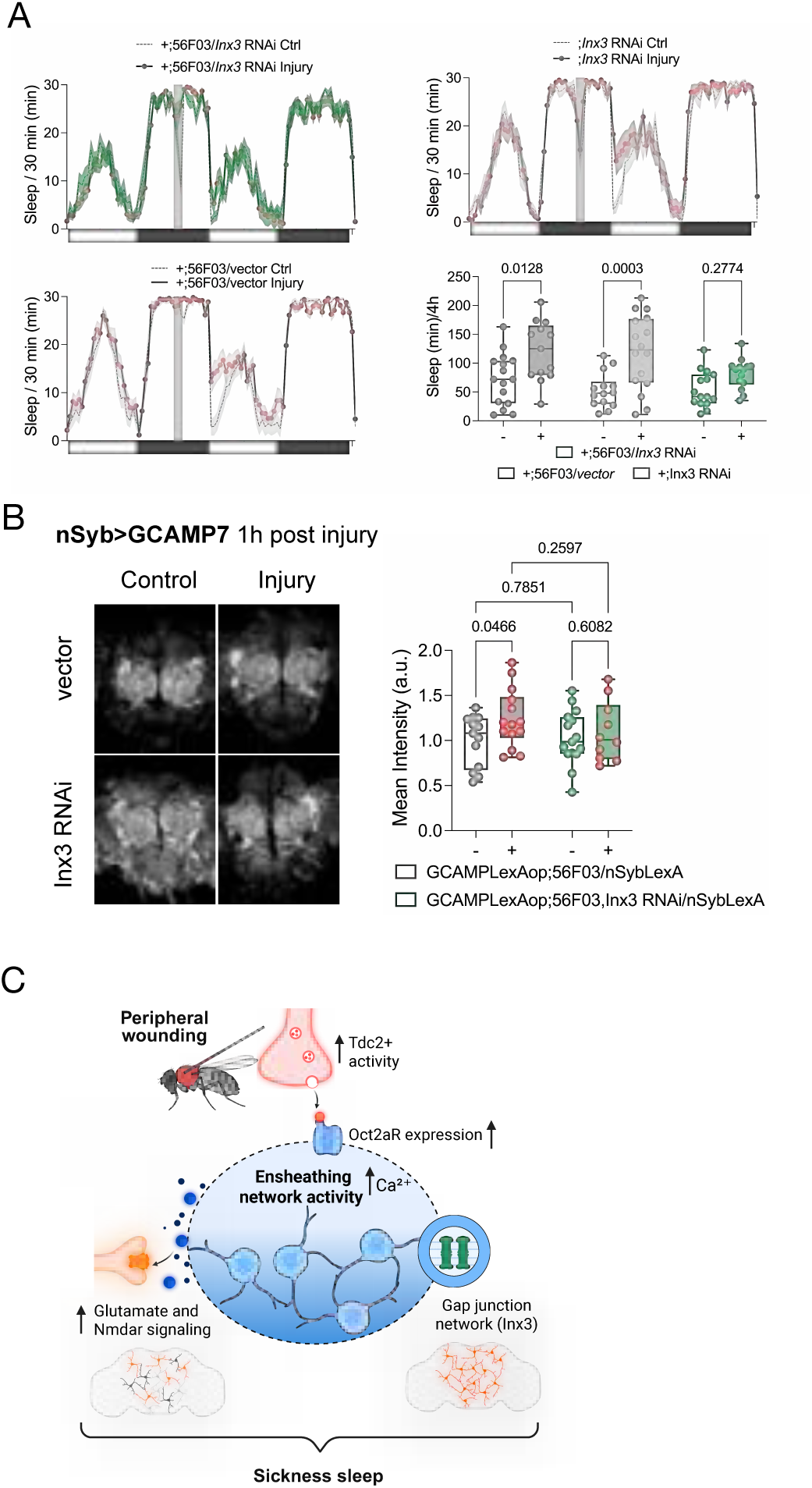
Glial network-dependent coordination of neuronal activity and sickness sleep. (A) Sleep profile after peripheral injection of flies with knockdown of Inx3 in ensheathing glia (green) and respective genetic controls (grey). (Bottom right) Quantification of time spent asleep between ZT0–4 for flies from L. ANOVA was done applying a Sidak multiple testing correction. For clarity, only test results for the control and injury comparison per genotype are shown. No statistical difference was found between the handled control flies of the different genotypes. n=16. (B) Ca signal in ensheathing glia 1h post injury, while driving the expression of the genetically encoded Ca indicator GCAMP7 with the nSyb-LexA driver in flies with Inx3 knockdown in ensheathing glia (green), compared to the genetic controls (grey). Data are normalized to the average of the control. Two-way ANOVAs followed by Tukey’s correction. n=5-8. (C) Model for glial coordination of sickness sleep. Peripheral wounding increases activity of octopaminergic (Tdc2+) neurons, which signal to ensheathing glia (EG) via the octopamine receptor Oct2aR. Oct2aR signaling modulates extracellular glutamate levels and neurons perceive this glutamate via ionotropic glutamate receptors, including NMDAR. Ensheathing glia activity increases brain-wide and a gap-junction-coupled network mediated by Innexin 3 facilitates neuronal brain-wide calcium increase. This EG-driven coordination of neuronal activity promotes sickness sleep, identifying ensheathing glia as active drivers — rather than passive bystanders — of the sickness response. Created in BioRender. Sehgal, A. (2026) https://BioRender.com/scfbktt

Together, these data support a model in which sickness involves distributed neuronal activation that relies on a gap junction dependent glial network (Fig. 5C).

## Discussion

Sickness is a conserved adaptive state in which animals reduce activity, alter sleep, and reallocate energy to promote survival and recovery (Davis and Raizen, 2017; Irwin, 2019; Kent et al., 1992). During sickness, peripheral immune signals communicate with the brain through circulating cytokines and vagal pathways (Watkins et al., 1995), and, recently, discrete neuronal populations in the mammalian brainstem and hypothalamus have been shown to control individual components of sickness behavior, such as feeding, activity or sleep (Darmohray et al., 2025; Ilanges et al., 2022; Jin et al., 2024; Osterhout et al., 2022). These studies have provided critical entry points into the neural control of sickness. However, they also raise a broader question that has remained largely unaddressed: how does the brain coordinate a global shift in neural state during sickness to modulate multiple behaviors through specialized circuits?

Here, we leveraged the cellular simplicity and genetic accessibility of the *Drosophila* brain to address this problem. Whole-brain single-cell profiling revealed that glial cells, rather than neurons, undergo the most extensive transcriptional remodeling following a peripheral, non-infectious injury. Although our single-cell profiling did not capture enough cells to resolve finer neuronal subclusters, this does not bear on the glial transcriptional changes that anchor our main findings. Functional analyses uncovered a glia-to-neuron signaling axis in which octopaminergic input activates a specific subtype of glia—ensheathing glia—to modulate glutamate homeostasis and promote sickness-associated sleep through ionotropic glutamate signaling.

Imaging experiments revealed that injury induces coordinated calcium dynamics in ensheathing glia and neurons across broad brain regions rather than within discrete loci. Notably, our calcium imaging was performed at a spatial and temporal resolution suited to capture brain-wide activation patterns rather than resolving individual calcium events at the cellular or subcellular level. Given that knocking down Innexin 3 specifically in ensheathing glia was sufficient to abolish sickness sleep, we suggest that the calcium signal is propagated through gap junctional coupling. Regardless, these findings position a glial network as a key substrate for coordinating global brain activity during sickness.

Our observation of widespread neuronal activation during sickness integrates independent studies in which specific behaviors were mapped to discrete loci, suggesting that local control nodes may operate within a broader, distributed framework that reorganizes brain activity globally. Neuromodulatory systems are well positioned to support such transitions, as they are known to regulate large-scale brain dynamics associated with arousal, attention, and sleep (Lee and Dan, 2012). The finding that octopamine, the fly ortholog of noradrenaline, and the glial *Octα2R* receptor are necessary nodes of sickness behavior and brain-wide activation, agrees with a model in which neuromodulatory inputs engage glial networks to broadcast state information across the brain, providing a substrate for coordinating neural activity beyond anatomically restricted circuits. The role of catecholamines in sleep modulation has recently been revised, shifting perspectives from the wake promoting properties of noradrenergic signaling (octopaminergic signaling in the fly) to a more nuanced framework. Infraslow NE oscillations modulate glymphatic clearance during sleep (Hauglund et al., 2025), participate in the buildup of sleep pressure to induce homeostatic sleep (Silverman et al., 2025), modulate NREM-REM cycles (Osorio-Forero et al., 2025) and alter sleep microarchitecture during stress (Antila et al., 2022). All in all, NE signaling might be critical not for sleep induction but rather promotion of sleep quality and restorative properties, and especially important in the context of sickness.

Ensheathing glia form continuous barriers around neuropils and axon tracts, placing them at strategic positions to influence large-scale brain activity (Kremer et al., 2017). However, while ensheathing glia have been implicated in several processes (Doherty et al., 2009; Otto et al., 2018; Stahl et al., 2018; Zhang et al., 2026) and even in the transmission of aversive information between sensory and high-order brain areas (Miyashita et al., 2023), the network role we describe here has not been previously shown. Notably, glutamate modulation is the common mechanism implicated in EG functions, and our data indicate that glutamatergic signaling similarly contributes to sickness sleep modulation, although parallel mechanisms of EG-neuron communication cannot be ruled out.

The EG gap junction protein Inx3 is also required for sickness sleep and the injury-induced rise in neuronal calcium, potentially serving as a substrate to broadcast signals across the brain. Circadian- and sleep-dependent regulation of innexin expression (Dopp et al., 2024) raises the possibility that the topology and permeability of these glial networks are dynamically tuned as in mammals (Cooper et al., 2025), allowing ensheathing glia to flexibly coordinate global brain states. Our data support this idea in the context of sickness behavior during which glial network integrity is essential for activating neurons across the brain and consolidating sickness sleep.

Together, our findings define a multi-step signaling cascade in which octopaminergic input engages ensheathing glial calcium dynamics, reshapes glutamate homeostasis, and promotes widespread neuronal activation to drive sickness-associated sleep. By positioning glial networks at the center of brain-wide state regulation, this work shifts the conceptual framework of sickness behavior away from strictly neuron-centric circuits and toward a distributed architecture in which glia integrate systemic cues and coordinate neural activity. This view offers a new lens through which to understand the neural basis of fatigue, malaise, and sleep alterations during illness, and suggests that targeting glial signaling pathways may provide novel strategies for modulating pathological sickness states.

## Material and Methods

### Fly rearing

Flies were raised at 25 °C and 60% relative humidity on fly food (including 64.7g/L cornmeal, 27.1g/L dry yeast, 8g/L agar, 61.6mL/L molasses, 10.2mL/L 20% tegosept, and 2.5mL/L propionic acid) under a 12–12 h light–dark cycle. All experiments were performed with female flies as male flies do not increase sleep robustly after injury. Experimental flies were collected at 2-3 days post eclosion, housed in cohorts of 30-40 flies and aged to 5-7 days.

For axenic fly generation (Zhang et al., 2025). Iso31 newborn fly embryos within 12 hours were rinsed in 100% ethanol, dechorionated in 10% bleach for 2 minutes, then immediately rinsed three times in germ-free PBS. The germ-free embryos were transferred to autoclaved standard germ-free molasses-cornmeal-yeast medium containing 1 mM kanamycin, 650 μM ampicillin (61-238-RH, MediaTech), and 650 μM doxycycline (D9891, Sigma-Aldrich). Germ-free flies were maintained on germ-free mediums with the three antibiotics for four generations and then maintained on the same medium without antibiotics for the next four generations.

For geneSwitch (inducible GAL4-UAS) experiments, food was prepared with either 500uM mifepristone (RU+ food) (Sigma-Aldrich M8046) or vehicle (equivalent volume of 100% EtOH). Flies were switched to RU+ at 2-3 days post-eclosion and allowed to eat for 3 days before behavioral testing.

For optogenetic experiments, flies were reared in constant darkness until eclosion. Posteclosion they were entrained to 12:12 LD cycles using dim red light. At 5-7 days flies were loaded into glass locomotor tubes containing 5% sucrose, 2% agar, and 1 mM all trans-retinal (ATR) or vehicle (equivalent volume of 100% EtOH). The end of the tube was covered in foil to prevent light from degrading the ATR. Illumination was achieved using LED strips fixed at intervals 8.75 cm apart to a 61 x 61 cm aluminum board. The board was positioned in the incubator on the shelf above the Drosophila Activity Monitors and positioned facing down on the shelf above the monitors. Upon loading, flies were illuminated with a 12:12 LD schedule of dim red light (400-700 lux). During GtACR1 silencing experiments, flies were exposed to 10s green light pulses (4000-6000 lux) for 10h starting after injury. Note that since injury is performed during the dark, the optogenetic inhibition expands to six hours of a dark background and four hours of dim red background.

### Fly stocks

The following lines are from the Sehgal lab stocks: CantonS Red, Iso31, sleepless P1 (backcrossed 5X to iso31), sleepless P1 (backcrossed 5X to iso31), fumin (backcrossed 5X to iso31), *nSyb*-GAL4, *repo*-GAL4, 58H05-GAL4, *nSyb*-GS-GAL4, *Daughterless-*GS, 56F03-GS-GAL4 (first used in this paper and tested by mCD8 expression after 72h 500mM RU feeding and available upon request).

The following lines were obtained from the Bloomington Drosophila Stock Center: 23E10–GAL4 (#49032), P{UAS-GtACR1.d.EYFP}attP2 (#92983), 56F03-GAL4 (#39157), 57C10-GAL4 (#39171), 57C10-LexA (#52817), *Tdc2*-GAL4 (#9313), 20XUAS-IVS-jGCaMP7b (#80907 and #79029), UAS-PV (#25030), 20XUAS-iGluSnFR (#59609), 3XLexAop-IVS-jGCaMP7b (#80915), UAS-R-GECO1 (#52222), UAS-*GLaz*-RNAi (#15389), UAS-*Octα2R*-RNAi (#50678), UAS-*Oct-TyrR*-RNAi(#28332), UAS-*Gdh*-RNAi (#51473), UAS-*Eaat1*-RNAi (#43287), UAS-*VGlut1*-RNAi (#40845), UAS-*Nmdar1*-RNAi (#25941), UAS-*Nmdar2*-RNAi (#26019), UAS-*Ekar*-RNAi (#28506), UAS-*ukar*-RNAi (#31991), UAS-*clumsy*-RNAi (#28351), UAS-*kaiR1D*-RNAi (#25852), UAS-*Grik*-RNAi (#25822), UAS-*GluRIB*-RNAi (#27673), UAS-*GluRIA*-RNAi (#27521), UAS-*mGluR*-RNAi (#25938), UAS-*mGluR*-RNAi (#34872), 14H09-AD;13B05-DBD (#86750), UAS-*Inx3*-RNAi (#30501), UAS-*Inx2*-RNAi (#29306), control line for TRiP RNAi lines Chr3 (#36303) and Chr2 (#36304).

The following lines were obtained from other sources: NP2222-GAL4 and *alrm*-GAL4 were shared by Marc Freeman, 9-137-GAL4 shared by R. Bainton, UAS-CaLexA (w; UAS-CD8::RFP, LexAop-CD8::GFP-2A-CD8::GFP; UAS-mLexA-VP16-NFAT, LexAop-CD2::GFP) shared by Jing W Wang, 84C10-AD; ChAT-DBD and Vglut-AD; 23E10-DBD shared by S. Dissel.

### Behavioral Assays

#### Peripheral Injury

Flies were removed from incubators at ZT18 and were typically exposed to a 1 hour light pulse during the protocol. This time corresponds to times when light pulses produce minimal phase shifts in locomotor activity rhythms (Suri et al., 1998). For injury, flies were removed from each glass activity tube and placed onto a CO^2^ pad. Injury was done by injecting the fly next to the humerus with a thin glass needle (Standard glass capillaries 1 mm O.D., WPI) pulled using a micropipette puller (Narishige) and attached to a syringe filled with PBS and 1% FCF (FCF Brilliant Blue Dye (FD&C blue no1 dye, Spectrum Chemicals). When the entire thorax turned blue and the injection medium was starting to get to the head, the needle was removed from the fly. Handled control flies were CO^2^ anesthetized for the same duration as the infected and injured groups (Kuo et al., 2012). For experiments involving sleep monitoring, flies were loaded into glass tubes containing 5% sucrose in 2% agar at 5-7 days post eclosion. On day three, individual flies were removed from the locomotor tubes and anesthetized. After injury, flies were placed back into glass tubes. When sleep measurement was not performed, flies were kept in regular food vials.

#### Sleep behavior

5-7 mated females were loaded into glass locomotor tubes containing 5% sucrose/2% agar food. Flies were given 1 d of acclimation before collection of data. Flies were maintained on a 12-h light/12-h dark cycle (except for optogenetic manipulations using GTACR which had a red/dark cycle and as indicated previously). Temperature was kept at 25 °C and 50% humidity. Sleep and locomotor activity data were collected in 1-min bins using the Drosophila Activity Monitoring (DAM) System (TriKinetics). Single beam monitoring was used except for experiments when Multibeam recording is explicitly mentioned. Sleep was defined as bouts of uninterrupted inactivity lasting for ≥5 min and analyzed with a custom software Insomniac3 (“Measuring Sleep in Drosophila | Springer Nature Experiments,” n.d.). For total sleep of day/night time sleep, data was analyzed for 24 or 12-h periods respectively. For sickness sleep, we used the total sleep minutes from ZT0 to ZT4 (4 h after lights on) after injury of both the injury and the handled control flies.

All sleep experiments were replicated two or three times with statistically significant results at least twice or three times respectively. Data shown in sleep plots are from a representative experiment.

### Quantitative PCR

RNAi efficiency was tested by qPCR using an ubiquitous GAL4 (daugtherless-G) to drive the expression of the RNAi targeting a larger cell population and avoid masking from expression in other cells types. We tested the four RNAi lines critical for the UAS-*Octα2R*-RNAi (#50678), UAS-*Nmdar1*-RNAi (#25941), UAS-*Nmdar2*-RNAi (#26019), UAS-*Inx3*-RNAi (#30501) that we considered critical for the project. All groups were treated for 72h with RU486 (Thermo Fisher Scientific) 500 mM in 5%sucrose 2% agar food. Flies were flash frozen within 2 h of Lights on. Heads were dissected and processed for RNA extraction using Trizol. In short, heads were hand held homogenized in 100ul of Trizol. 400ul extra Trizol was added followed by 100 ul chloroform, vortexed, and wait 10 min room temperature (RT). Aqueous phase collected and mixed with one volume of isopropanol. After 10 min at RT samples were spined down, isopropanol removed and add 100ul 70% ethanol, spined down again and pellet air dried. DNA was removed using the DNA-free™ DNA Removal Kit (Fisher) according to manufacturer’s instructions. 1000ug of RNA was used for cDNA conversion using High-Capacity cDNA Reverse Transcription Kit (LifeTech) according to manufacturer’s instructions. Quantitative real-time PCR (RT-qPCR) was performed using PowerUp™ SYBR™ Green Master Mix (Applied Biosystems) on a QuantStudio™ Real-Time PCR System (Applied Biosystems).

### Primers

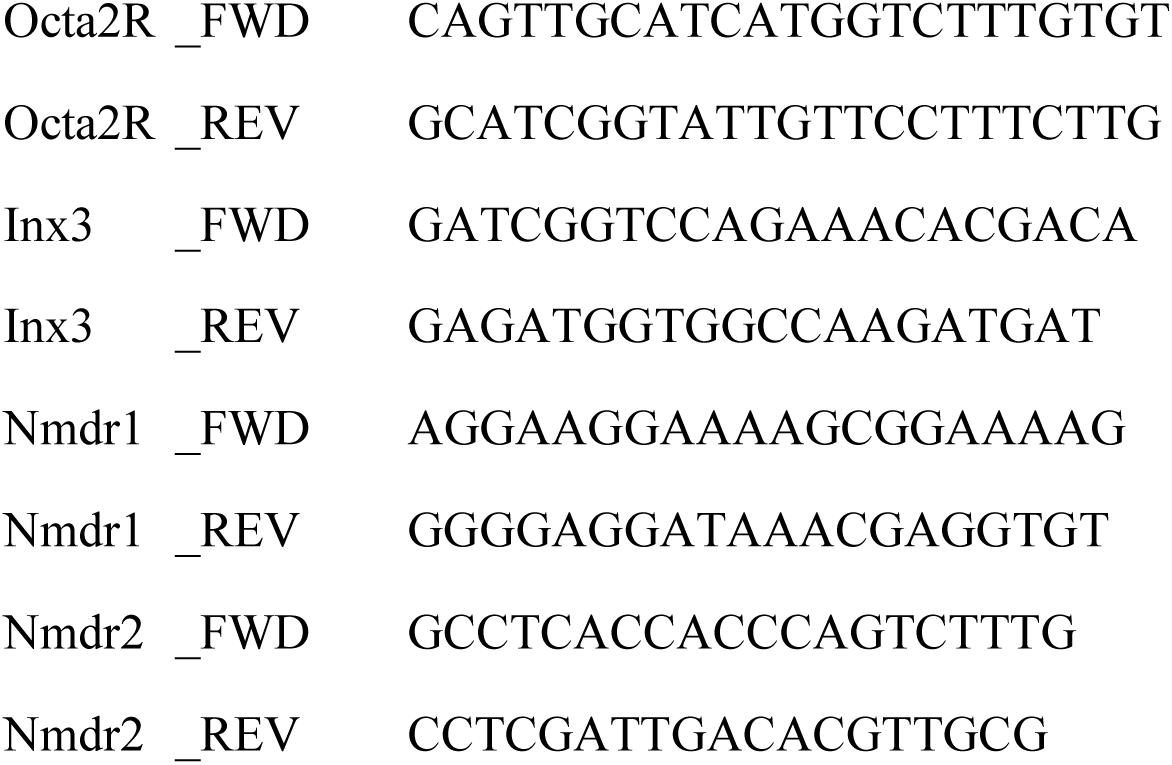

### scRNA-seq sample preparation and 10xGenomics sequencing

100 female CantonS red flies were dissected in ice cold Schneider’s medium. Optic lobes were removed and transferred to a tube with 3mg/ml of dispase. Samples were incubated in a thermoshaker for 15 min at 25C and 1000 rpm. To stop the enzymatic reaction, cells were rinsed three times with Schneider’s with table top spins in between. Cells were resuspended in 60 ul of DPBS with 0.04% BSA. Cell suspensions were FACS sorted using an Aria Cell Sorter (using a 100 um nozzle and 1.25 drop precision) to ensure cell viability and adjust concentration. Cells were kept on ice until processed for 10x Genomics. Single Cell 3’ Library and Gel Bead Kit following the manufacturer’s recommendation protocol and the libraries were sequenced on an Illumina NovaSeq 6000 in the Centre for Applied Genomics at the Children’s Hospital of Philadelphia (CHOP).

### scRNA-Seq analysis using SCANPY

We aligned and quantified the sequencing reads using the CellRanger (v7.2.0) count command. We captured an estimated 4546 cells in the control sample, with a mean reads per cell of 54269, and an estimated 4203 cells in the injury sample, with a mean reads per cell of 50309. We then performed downstream analysis on the output filtered feature barcode matrices using the Python package SCANPY (v1.9.6).

For the preprocessing of the data, we first filtered out outlier cells and cells with mitochondrial percentage over 20% (damaged/dying cells). Then we filtered ambient RNA with SoupX (from ∼13000 genes to ∼8000 genes), and filtered doublet cells with scDblFinder (∼6% cells called doublets). We integrated filtered control and injury data sets with scVI, taking into account batch effects for better mixing. The resulting merged data set contains 4201 cells from the control group and 3713 cells from the injury group. We then normalized the read counts using shifted logarithm with Scran estimated size factors. Finally, we performed principal component analysis (PCA) on the top 4000 highly variable genes, and the variance ratio levelled off at around 50 PCs. We created umap and tsne plots to visualize the data set.

We clustered the scVI integrated data using the leiden algorithm (scanpy.tl.leiden), using the top 50 principal components, at various resolutions ranging from 0.5 to 20. For the automated annotation of the cells, we used two methods, scArches (scVI tools v1.0.4) and CellTypist (v1.6.2). With scArches, we integrated our data set with a reference data set (ref. Davie et al 2018), and then used the k-nearest neighbor algorithm to transfer the cell type labels from the reference data set to our data set. Celltypist uses a logistic regression model trained on the reference data set to predict the cell type for each cell in our data set based on its transcriptomic profile. For both methods, we marked cells annotated with less than 80% certainty as unknown. We then took the combined annotation from these two methods, going with CellTypist for the few cells where they disagree since its annotation aligns slightly better with expression of known marker genes. Finally, we manually annotated 13 leiden clusters from resolutions 0.5, 5, 15 and 20 according to the results of this combined automated annotation. Since some of the neuronal clusters are too small for downstream differential expression analysis, we also annotated the *nSyb* positive cells with more general neuronal cell types. Using the four neuronal markers (Gad1 for gabaergic cells, VGlut for glutamatergic cells, VAChT for cholinergic cells and Vmat for monoaminergic cells), we annotated the neurons with their most highly expressed marker.

Differential Expression analysis on each of the large cell types was performed using the pseudobulk method. We constructed 3 pseudobulk samples for each cell type and each condition by randomly dividing the cells into 3 samples and aggregating the gene counts. We then ran DESeq2 on the pseudobulk data to generate our list of differentially expressed genes for each cell type.

### Immunostaining

Adult fly brains were dissected in cold phosphate buffered saline (PBS) and then fixed in 4% paraformaldehyde for 20 minutes at room temperature. Brains were then washed twice for 15 minutes in 1 ml PBS with 0.3% Triton-X (PBST) and blocked for 1h in 10% Normal Goat Serum in 0.3% Triton-X (PBST). We performed immunostaining for CaLexa using antibodies against GFP (1:500; rabbit) and RFP (1:500; rat, chromotek, 5f8-100). Primary and secondary antibody incubations were performed in PBS + 0.3% Triton X-100 (PBST). Following primary antibody staining, brains were washed twice for 15 min in 1 ml PBS with 0.3% Triton-X (PBST), stained for 2 h with Alexa 488 anti-rabbit and Alexa 647 anti-rat (1:500; Invitrogen A-11008, Invitrogen A-21247), washed twice for 15 min in 1 ml PBS with 0.3% Triton-X (PBST), mounted and imaged. We also repeated the CaLexA experiments visualizing the endogenous signal with similar results. Images presented in the paper correspond to the antibody conditions. Samples were then placed in 50% glycerol and mounted in Vectashield: H1000. Control and injury brains were mounted on the same slide and imaged using a Leica Stellaris 8 confocal microscope using a 20x objective. Sum projections were used to measure GFP and RFP intensities in specific ROIs as indicated in each experiment. Data are presented as GFP:RFP ratios normalized to the control group. Images correspond to max z-projections for clarity using the Fire LUT. All images are available upon request.

### Ex vivo brain imaging

Adult fly brains were dissected in imaging buffer (NaCl 108mM, KCl 5mM, CaCl2 2 mM, MgCl2.6H2O 8.2mM, NaHCO3 4mM, NaH2PO4.H2O 1mM, D-glucose 10mM, Na.HEPESbase 9mM HEPES acid 11mM, adjusted to pH 7.3) and mounted in poli-D-Lysine coated glass dishes with 70 μl imaging buffer (MATTEK, P35G-1.5-10-C). 5 min after mounting, brains were imaged with a Leica Stellaris 8 confocal microscope with a 10× objective, 488 nm (for GCAMP and iGkusnFR) iand 561 nm (for GECO1) laser excitation. Brains were imaged in sets of 4 every 10 s for the indicated time. For one time point experiments, three imaging rounds were performed and average fluorescence was used for data presentation. For experiments representing time courses, 3 base line acquisitions were done before any drug application. Drugs and neurotransmitters were dissolved in the imaging buffer in a 2x concentration. Equal volume to the initial incubation buffer was applied by carefully pipetting the solution over the brains. For instances when brains were moved during the drug application, data was excluded. Drugs and neurotransmitters: Tetrodotoxin final concentration 1mM, TTX (Fisher, NC1588755), Tyramine (Sigma, T2879), Octopamine (Sigma, O0250). For data analysis, sum intensity of the z-protections of each imaging cycle were used. Optic lobes were excluded.

### Long term brain in vivo preparation

3-5 day old female flies were used. Opening of the head cuticle was performed with slight modifications to Huang et al., 2018. In brief, flies were cold anesthetized and mounted in a custom-made silicon fixture with a fused silica optical fiber that was placed in between the head and thorax using UV Epoxy (Norland Optical Adhesive NOA 68). To create the optical window, we used 0.001mm Tip Tungsten Needles (Fisher, NC9221118) under a humidified ambient using a 5x magnifying scope. Cuticles, air sacs and fat bodies were removed until the brain was visible. Immediately after surgery, we applied UV epoxy (Norland Optical Adhesive NOA 68, Norland, 36-427; 1.54 refractive index; ∼99% optical transmission for wavelengths between 420 and 1000 nm) and cured it. After a few minutes, flies were released from the silicone fixture and returned to vials containing complete food (see Fly husbandry). Flies were allowed to recover for at least 48h and were used for imaging up to 7 days post surgery.

### 2 photon imaging

Flies were mounted on custom-made silicon fixtures fixing the thorax of the fly to an optic fiber using UV Epoxy (Norland Optical Adhesive NOA 81, Norland, 36-428). A small piece of glass coverslip No.1 was glued to the optical window using NOA 68. Flies were allowed to recover for 10 min before imaging. Flies were injected or lightly manipulated (handled control) and immediately placed under the imaging scope. Images were taken using a Bruker nanosystems Ultima two-photon microscope equipped with a Chameleon ULTRA I laser directed through a resonant scanning galvanometer with a 20X Olympus objective at a digital zoom of 2X. Images were acquired at a resolution of 256 x 256 pixels in Z-stacks slices 15 μm apart. Image stacks were acquired for 5 min every fifteen minutes for three hours after injury. For the subsequent three hours images were taken every 30 min. Only flies that survived the imaging round and still responded vividly to tactile stimulus were included in the analysis.

For each brain, the image stack acquired during the 5 min acquisition was converted into a Z-sum projection to draw an ROI around the signal. An ROI was drawn outside to account for background. The mean pixel intensity for this area was computed for each Z projection and the signal from the background ROI subtracted. For each time point, intensities of the 5 min recording were averaged prior exclusion of the first minute to avoid potential artifacts, every fly time course was scaled between 0 and 1 and further normalized to F_0_ where F_0_ is the pre injury image (ΔF= (F-F_0_)/ F_0_).

## Statistical Analysis

All tests were performed using GraphPad Prism 10. Statistical parameters are listed in the caption of each figure. In brief, in sleep experiments, for 30 minute binned profiles, data is shown representing means and SEMs. For box and quartile plots, data is shown representing medians and displaying every data point. For ANOVAs, Holm-Sidak or Sidak multiple testing corrections were performed depending on the pre-selection of comparisons. In most graphs only comparisons between the control and injury case are shown and the entire statistic tree is included in Table S2.

### Limitations of the study

The findings in this body of work should be considered alongside the following limitations. Our single-cell profiling did not capture enough cells to resolve finer neuronal subclusters; however, this does not bear on the glial transcriptional changes that anchor our main findings. Similarly, our calcium imaging was performed at a spatial and temporal resolution suited to capture brain-wide activation patterns rather than resolving individual calcium events at the cellular or subcellular level. Therefore, the Inx3-dependent gap junction coupling role in the propagation of signals should be considered as suggestive and hopefully the new techniques and approaches currently in development in the field will allow for a more detailed description in future studies.

## Data availability

scRNAse of Drosophila brain. GEO Series GSE333749

## Supplementary Data

Table S1. Differential Expression Gene Analysis. Tab 1, all genes and cells. Tab 2 differentially expressed. Tab 3, Octα2R across cell types.

Table S2. Differential expression of EG subclusters.

## Acknowledgements

We thank the Centre for Applied Genomics Team for performing 10x Genomics and Jennifer Jakuvowski for performing FACS-based cell isolation and flow cytometry analysis in the Penn Cytomics and Cell Sorting Shared Resource Laboratory. We thank Gregory R Grant and Antonijo Mrčela for advice and support for single cell sequencing data analysis. We thank Daniel Iasconne Paula Haynes and Erick Astacio for advice guidance during the ex vivo brain imaging.

We thank Cheng Huang and Ruibao Zhu for teaching and guidance for the long term in vivo imaging. We thank M. Freeman, R. Bainton, and S. Dissel for sharing fly lines. This work was supported by SNSF Postdoc.Mobility Fellowship.

---

**Figure S1.**
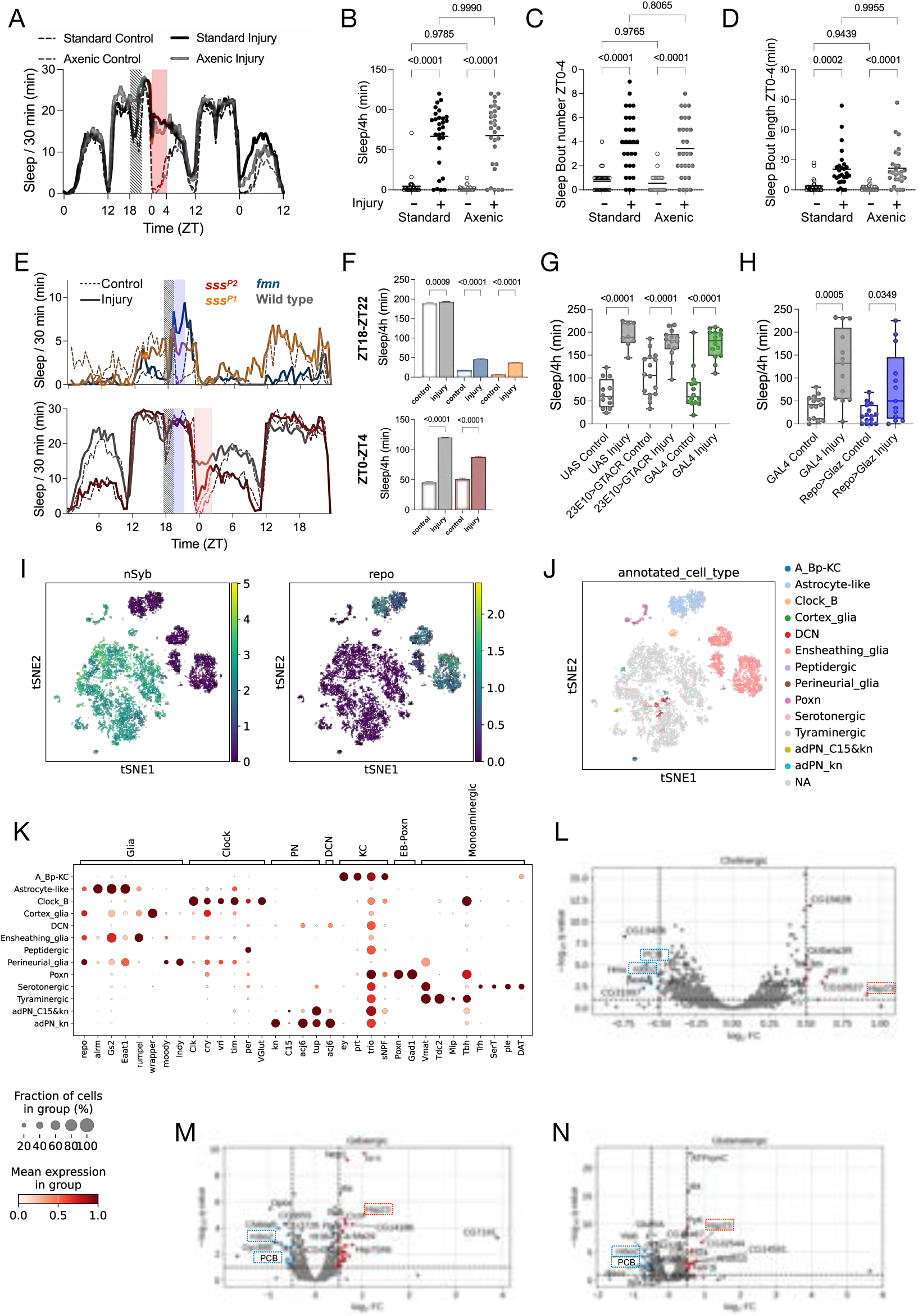
Sickness sleep is independent of the microbiome and does not appear to use mechanisms of daily sleep (A) Sleep profile (average time spent sleeping in consecutive 30-min segments; mean) of flies after peripheral injection in standard or axenic (sterile) conditions. Dashed lines correspond to handled control flies, thick lines to injured flies. Grey vertical bar indicates injection time. Red area highlights the time window when sickness sleep is observed and used to quantify sleep and bouts in B-D. (B–D) Quantification of sleep (B) and sleep bout length (C) and number (D) for flies from A between ZT0 and ZT4. Two-way ANOVAs followed Tukey’s correction. n=32 flies per group. (E) Sleep profile of flies after peripheral injection for three sleep mutants *sleepless P1* (*sss^p1^*), *sleepless P2* (*sss^p2^*), *fumin* (*fmn*) and the wild type control in the same genetic background (iso31). Genotypes are split into top (*sss^p2^* and wild type) and bottom (*sss^p1^* and *fmn*) for better visualisation of sickness sleep in the low sleepers *sss^p1^* and *fmn*. n=16. (F) Quantification of minutes of sleep between ZT18-ZT22 (top, for low sleeping mutants *sss^p1^* and *fmn*) and ZT0–4 (bottom, for wild type and *sss^P2^*)) from E. Two-way ANOVA was done while specifying the comparisons between control and injury for each genotype; therefore Sidak multiple testing correction was applied. n=16. (G and H) Quantification of sleep after injury while optogenetically inhibiting sleep promoting cells using the 23E10-GAL4 driver (G) and after knockdown of the lipid transporter *Glaz* in glia (H). In G all groups were treated with ATR food for two days before injury. Two-way ANOVAs followed Tukey’s correction. n=8-16. (I) tSNE of 7k cells characterized by single-cell RNA sequencing of 5-day-old fly brains (controls and injury). Colors indicate the expression of the neuronal marker nSyb (left) and the glia marker repo (right). Scale units are log1p normalized expression. (J) tSNE displaying cell types from iterative Leiden clustering and multi-step annotation. 3,094 colored cells were assigned to 13 clusters spanning four glial types and several neuronal classes. See S1K for marked gene expression. (K) Marker gene expression across the clusters annotated in J. (L-N) Volcano plots from DEG in in cholinergic cells (L), gabaergic cells (M) and glutamatergic cells (N). DEGs are highlighted in blue (downregulated) and red (upregulated) and include genes with fdr < 0.05 and log fold change lower than −0.5 or greater than 0.5. Annotated genes correspond to the top 10 DEG. Dotted squares indicate the three genes (PCB, robo2 and Hsp23) with the same trend in the three clusters.

**Figure S2:**
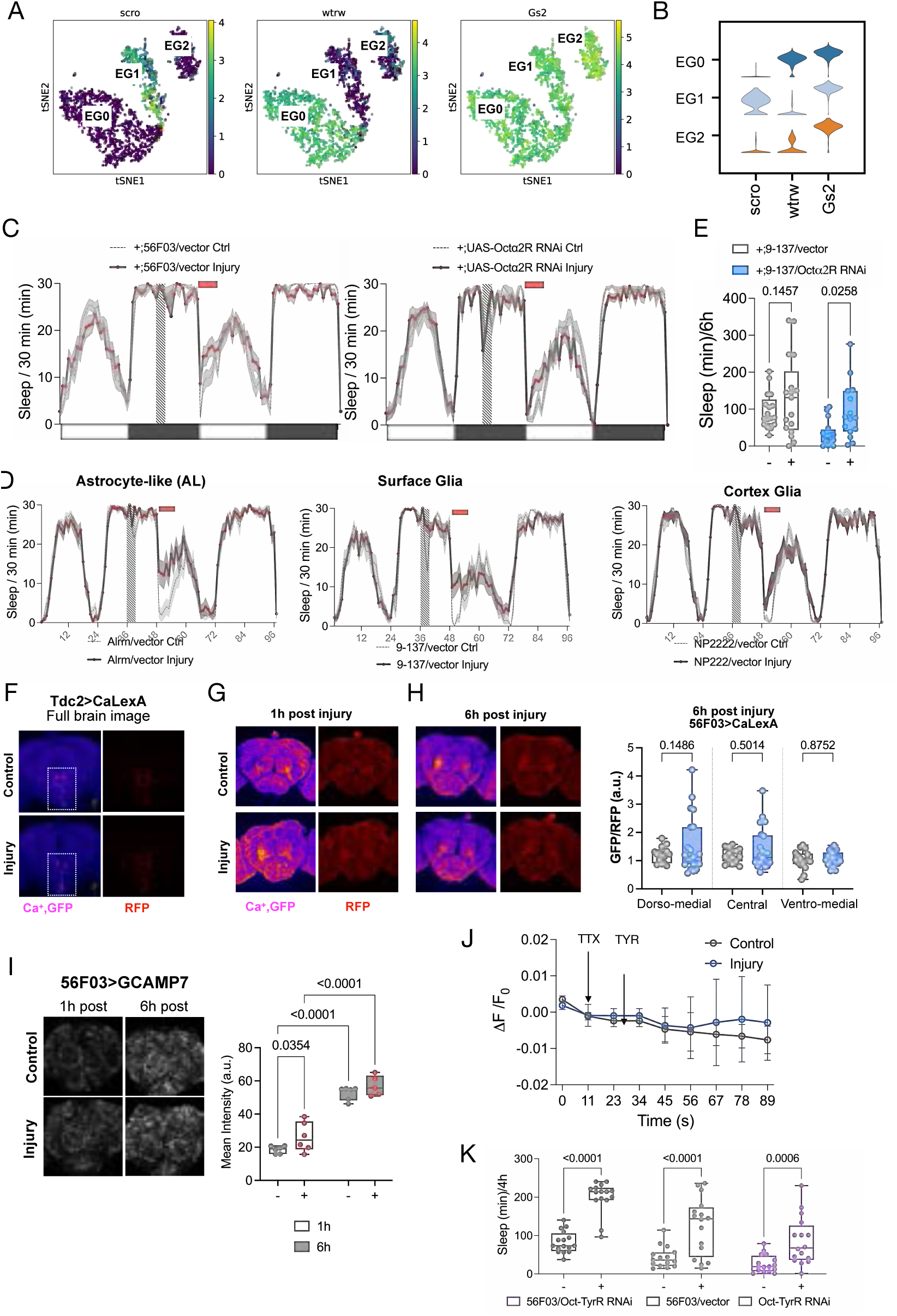
Octopamine signals through ensheathing glia to regulate sickness sleep (A and B) tSNE (A) and violin plots (B) of the three EG sub clusters characterized by expression of scro (EG1), wtrw (EG0) and absence of both (EG2). (C) Sleep profiles complementary to Figure 2A representing the GAL4 and UAS control genotypes. (D) Sleep profiles complementary to Figure 2C representing the GAL4 genotypes. UAS control genotypes were not assayed in this experiment since they were redundant with those in 2A and S2C. (E) Quantification of time spent asleep between ZT0–6 for flies with surface glia knockdown of Octa2R (from 2C). ANOVA with Sidak multiple testing correction. Only comparison between control and injury is shown for better visualization (NOTE handled controls are significant only when quantified from 0-4, and are shown in Figure 2D). n=16. (F) Full brain image from 2G displaying the calcium signal in Tdc2 cells expressing CaLexA (pseudocolored using the Fire LUT from ImageJ) and an internal control (red). Square highlights the ventral cells shown in 2G. (G) Ca imaging and quantification of ensheathing glia, while driving the expression of CaLexa with the 56F03-GAL4 driver. RFP internal control of 1h post injury is shown in 2G. Quantifications shown in 2H. ANOVA with Holm-Sidak multiple correction. n=10-16. (H) Ca imaging and quantification of ensheathing glia 6h post injury, while driving the expression of CaLexA with the 56F03-GAL4 driver. For analysis, brain was subdivided into PI region, central brain and ventral brain. Data are normalized to the average of the control brain data for each region. ANOVA with Holm-Sidak multiple correction. n=10-16. (I) Representative images and quantification of Ca in ensheathing glia 1h and 6h post injury using the genetically encoded Ca indicator GCAMP7 with the 56F03-GAL4 driver. ANOVA was done applying a Sidak multiple testing correction specifying the time matched comparisons. n=5-8. (J) Ca levels in EG after bath application of 2.5 mM of tyramine (TYR). Brains were dissected 1h after injury and imaged in a saline bath containing 1uM TTX to block neuronal activity. ANOVA with Sidak multiple correction. (K) Sleep profiles and quantification of time spent asleep between ZT0–4 of adult-specific knockdown of the Tyramine receptor (Oct-TyrR) in ensheathing glia (56F03-GAL4). Sleep profile not show due to space managing but available upon request. ANOVA was done applying a Holm–Sidak multiple testing correction.

**Figure S3.**
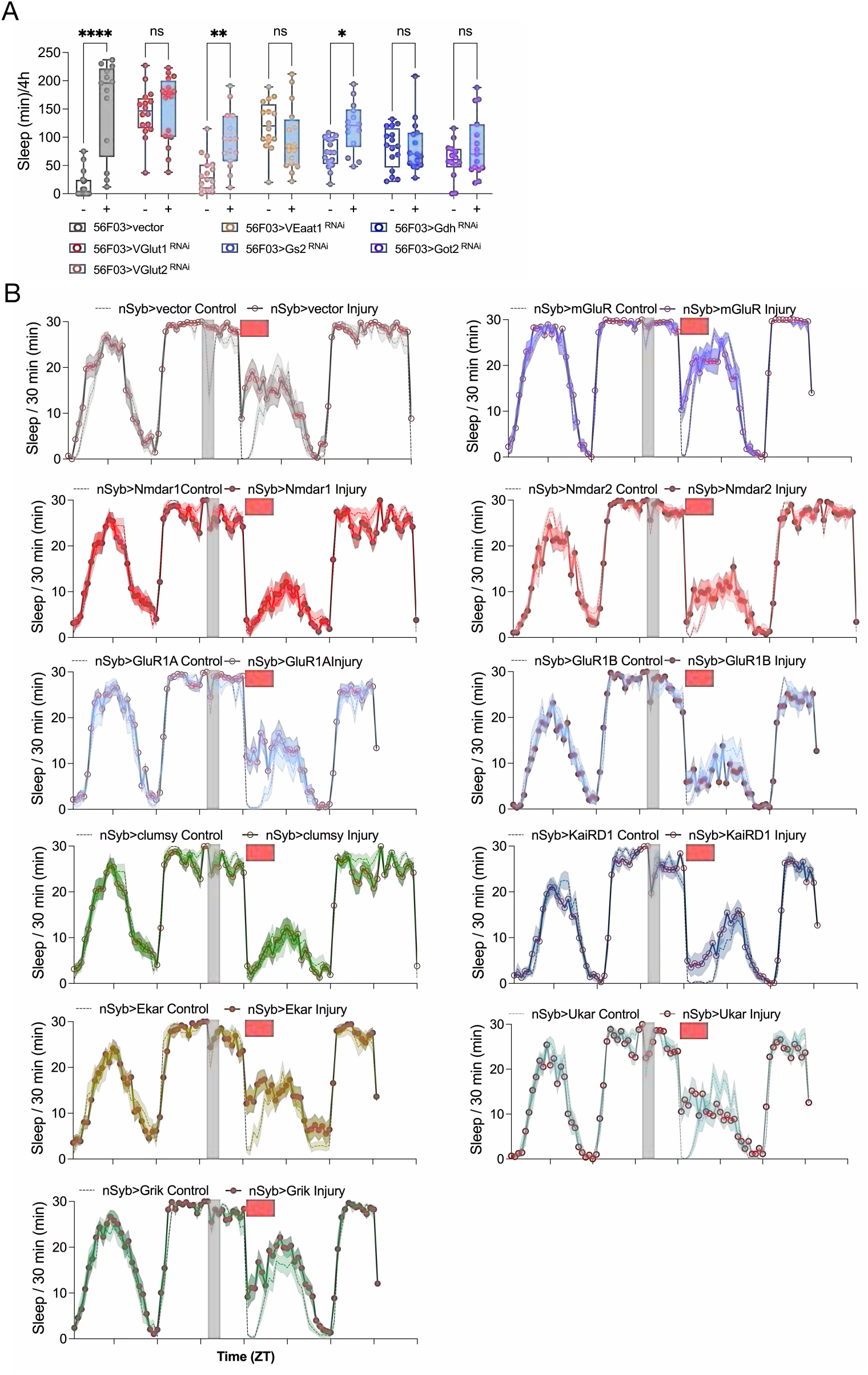
(A) Quantification of time spent asleep between ZT0–4 by flies with knockdown of glutamate transporters and metabolism genes in ensheathing glia (56F03-GAL4). ANOVA was done specifying comparisons between control and injury for each genotype; therefore Sidak multiple testing correction was applied. n=16,*<0.05,**<0.01,****<0.0001. (B) Sleep profiles corresponding to Figure 2C, regarding knockdown of glutamate receptors in neurons (nSyb-GAL4).

**Figure S4.**
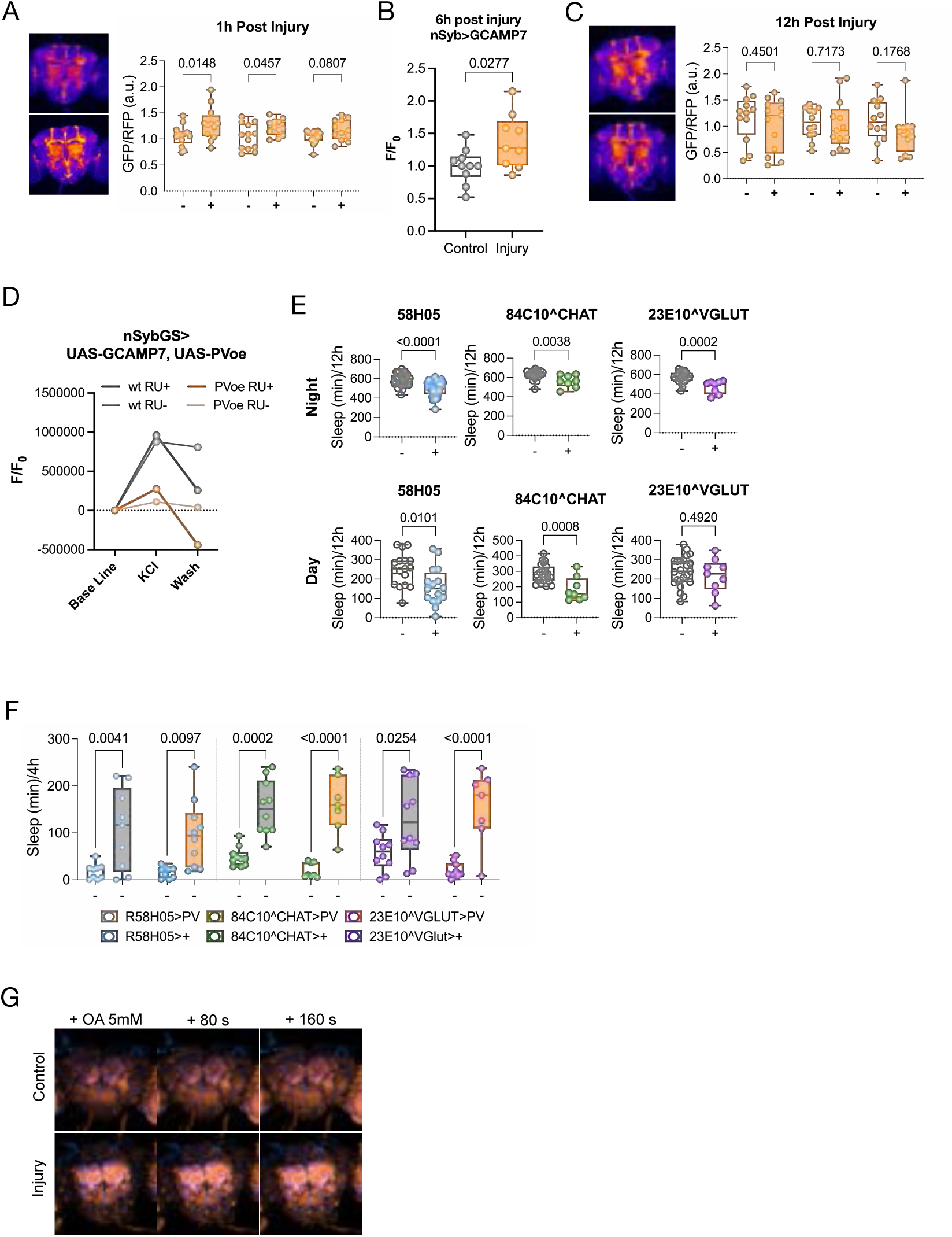
Dissecting the role of calcium signaling in sickness sleep (A) Ca imaging and qualification of neurons 1h post injury, while driving the expression of CaLexA with the nSyb-GAL4 driver. For analysis, brain was subdivided into a PI region, central brain and ventral brain (see diagram). ANOVA with Holm-Sidak multiple correction. n=10-16. (B) *In vivo* 2P imaging of Ca using GCAMP7 in neurons. Optical windows were opened at 6h post injury and brains were immediately imaged. n=9-10. (A) Ca imaging and qualification of neurons 12h post injury, while driving the expression of CaLexa with the nSyb-GAL4 driver. For analysis, brain was subdivided into a PI region, central brain and ventral brain (see diagram). ANOVA with Holm-Sidak multiple correction. n=10-16. (D) Ex vivo imaging of brains with conditional expression of the Calcium buffer protein Parvalbumin in neurons. Expression of the transgene on its own already reduced the response to KCl measured by intracellular calcium changes (imaged with the genetically encoded indicator GCAMP7). The application of RU486 (RU+), further reduced intracellular calcium after extracellular media dilution (wash). (E) Quantification of time spent asleep during the day and night of flies expressing PV in sleep promoting centers ellipsoid body (58H05-GAL4), dorsal fan body (58H05^ChAT and 23E10^VGlut). Umpaired t-test. n=10-16. Sleep profiles available upon request. (F) Quantification of time spent asleep between ZT0–4 of flies expressing PV in sleep promoting centers ellipsoid body (58H05-GAL4), dorsal fan body (58H05^ChAT and 23E10^VGlut). ANOVA was done specifying the comparisons between control and injury for each genotype; therefore Sidak multiple testing correction was applied. n=10-16. Sleep profiles available upon request. (H) Representative images of flies expressing Ca indicators in neurons (nSyb-LexA>LexAop-GCAMP7, orange) and EG (56F03-Gal4>UAS-GECO1, blue).

**Figure S5.**
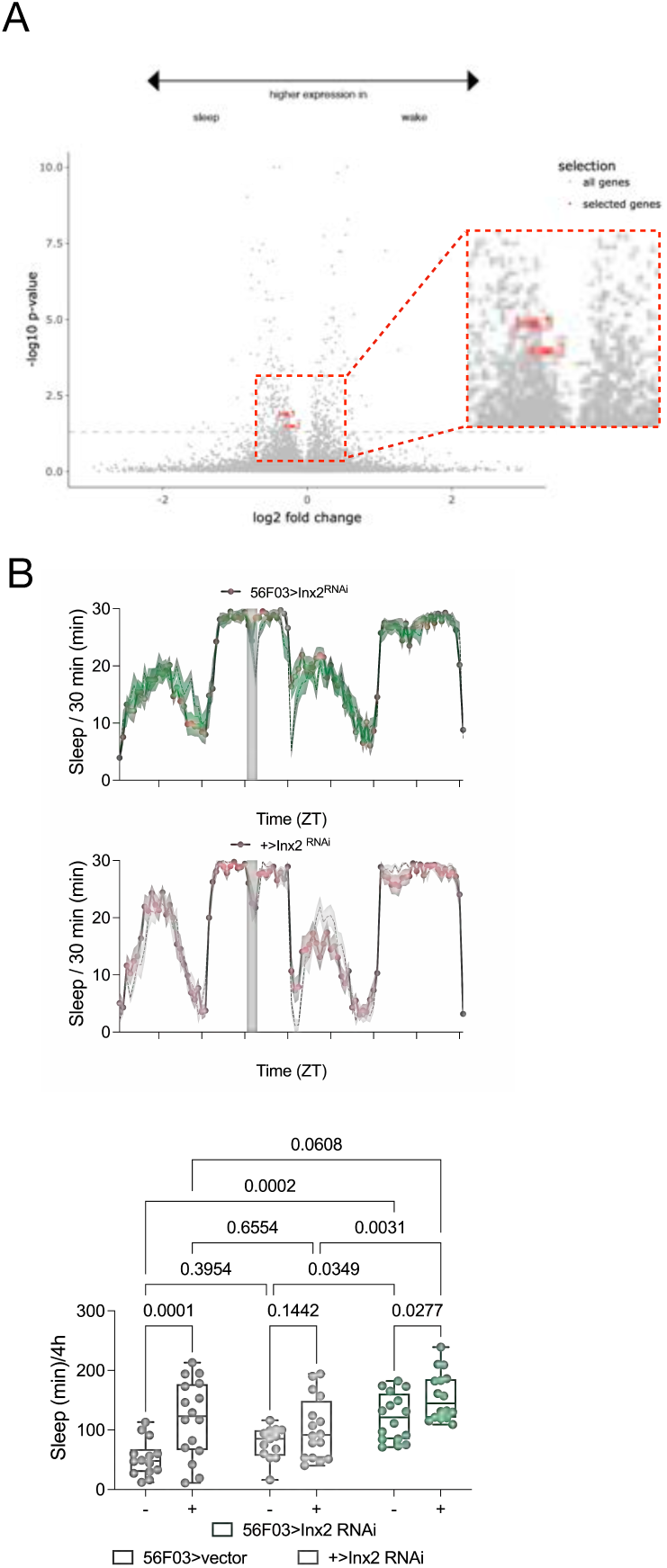
Role of Innexins in sickness sleep (A) Volcano plot representing the expression of Inx2 and Inx3 in the EG_1 cluster from Dopp et al. Genes on the left increase in sleep vs wake. Plot adapted from https://joana-dopp.shinyapps.io/Fly_Sleep_Single_Cell_v1/. (B) Sleep profile after peripheral injection of flies with knockdown of Inx2 in ensheathing glia (green) and respective genetic controls (grey). (B) Quantification of time spent asleep between ZT0–4 for flies from L. Note the 56F03>vector is the same as in Fig. 5A. ANOVA was done applying a Sidak multiple testing correction. For clarity, only test results for the control and injury comparison per genotype are shown. n=16.

**Figure S6.**
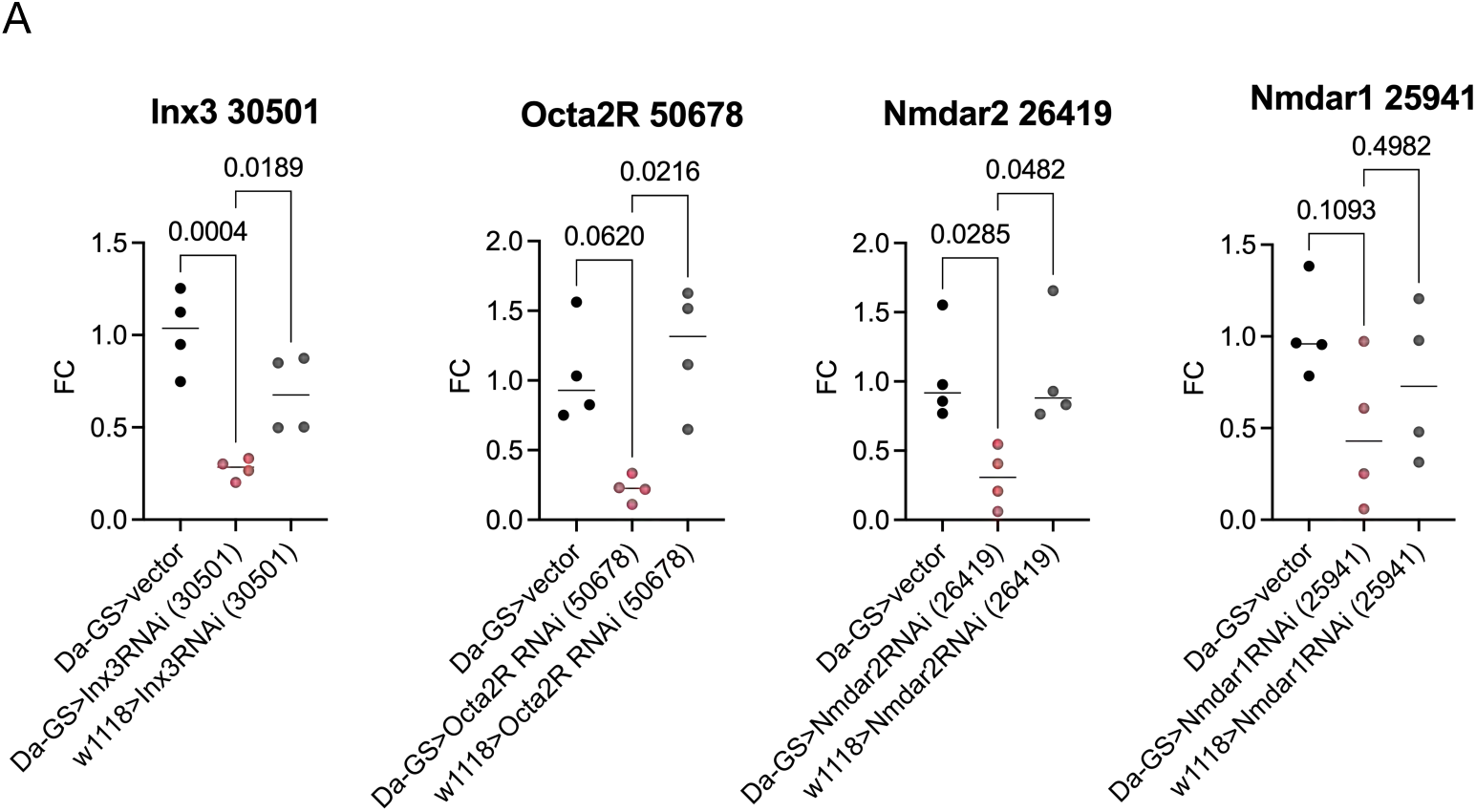
Analysis of RNAi efficacy (A) Validation of RNAi knockdown efficiency by RT-qPCR. Relative transcript levels of the target gene were quantified by RT-qPCR in flies expressing each of the three RNAi lines; UAS-Octα2R-RNAi (#50678), UAS-Nmdar1-RNAi (#25941), UAS-Nmdar2-RNAi (#26019), UAS-Inx3-RNAi (#30501). Expression was normalized to the reference gene Rpl32. Fold change (FC) is shown relative to the GAL4 > attP control. RNAi parental lines crossed to the GAL4 genetic background strain were included as additional controls. For Nmdar1 data shows a clear trend of reduced expression in the experimental compared to the GAL4 and UAS control despite not reaching significance. Statistical significance was assessed by one-way ANOVA with Dunn’s multiple comparisons test, n=4.

## References

Allen, N.J., Lyons, D.A., 2018. Glia as architects of central nervous system formation and function. Science 362, 181–185. 10.1126/science.aat0473

Antila, H., Kwak, I., Choi, A., Pisciotti, A., Covarrubias, I., Baik, J., Eisch, A., Beier, K., Thomas, S., Weber, F., Chung, S., 2022. A noradrenergic-hypothalamic neural substrate for stress-induced sleep disturbances. Proc Natl Acad Sci U S A 119, e2123528119. 10.1073/pnas.2123528119

Bittern, J., Pogodalla, N., Ohm, H., Brüser, L., Kottmeier, R., Schirmeier, S., Klämbt, C., 2021. Neuron–glia interaction in the Drosophila nervous system. Developmental Neurobiology 81, 438–452. 10.1002/dneu.22737

Blum, I.D., Keleş, M.F., Baz, E.-S., Han, E., Park, K., Luu, S., Issa, H., Brown, M., Ho, M.C.W., Tabuchi, M., Liu, S., Wu, M.N., 2021. Astroglial Calcium Signaling Encodes Sleep Need in Drosophila. Curr Biol 31, 150–162.e7. 10.1016/j.cub.2020.10.012

Cahill, M.K., Collard, M., Tse, V., Reitman, M.E., Etchenique, R., Kirst, C., Poskanzer, K.E., 2024. Network-level encoding of local neurotransmitters in cortical astrocytes. Nature 629, 146–153. 10.1038/s41586-024-07311-5

Černe, U., Horvat, A., Sanjković, E., Kozoderc, N., Kreft, M., Zorec, R., Scholz, N., Vardjan, N., 2025. Ca2+ excitability of glia to neuromodulator octopamine in Drosophila living brain is greater than that of neurons. Acta Physiol (Oxf) 241, e14270. 10.1111/apha.14270

Charles, A., 1998. Intercellular calcium waves in glia. Glia 24, 39–49. 10.1002/(sici)1098-1136(199809)24:1%3C39::aid-glia5%3E3.0.co;2-w

Chen, A.B., Duque, M., Rymbek, A., Dhanasekar, M., Wang, V.M., Mi, X., Tocquer, L., Narayan, S., Legorreta, E.M., Eddison, M., Yu, G., Wyart, C., Prober, D.A., Engert, F., Ahrens, M.B., 2025. Norepinephrine changes behavioral state through astroglial purinergic signaling. Science 388, 769–775. 10.1126/science.adq5233

Chever, O., Dossi, E., Pannasch, U., Derangeon, M., Rouach, N., 2016. Astroglial networks promote neuronal coordination. Sci Signal 9, ra6. 10.1126/scisignal.aad3066

Cooper, M.L., Selles, M.C., Cammer, M., Gildea, H.K., Sall, J., Chiurri, K.E., Saab, A.S., Liddelow, S.A., Chao, M.V., 2025. Astrocytes connect specific brain regions through plastic gap junctional networks. bioRxiv 2025.07.18.665573. 10.1101/2025.07.18.665573

Cornell-Bell, A.H., Finkbeiner, S.M., Cooper, M.S., Smith, S.J., 1990. Glutamate induces calcium waves in cultured astrocytes: long-range glial signaling. Science 247, 470–473. 10.1126/science.1967852

Cuddapah, V.A., Hsu, C.T., Valle Sirias, F., Li, Y., Shah, H.M., Saul, C., Killiany, S., Guevara, C., Shon, J., Yue, Z., Gionet, G.L., Putt, M.E., Sehgal, A., 2025. Sleep drive, not total sleep amount, increases seizure risk. Nat Commun 16, 6967. 10.1038/s41467-025-62311-x

Darmohray, D., Sima, J., Chen, C.-H., Silverman, D., Chen, C., Xu, A., Yao, Y., Dan, Y., 2025. Brainstem circuit for sickness-induced sleep. Sci Adv 11, eady0245. 10.1126/sciadv.ady0245

Davie, K., Janssens, J., Koldere, D., De Waegeneer, M., Pech, U., Kreft, Ł., Aibar, S., Makhzami, S., Christiaens, V., Bravo González-Blas, C., Poovathingal, S., Hulselmans, G., Spanier, K.I., Moerman, T., Vanspauwen, B., Geurs, S., Voet, T., Lammertyn, J., Thienpont, B., Liu, S., Konstantinides, N., Fiers, M., Verstreken, P., Aerts, S., 2018. A Single-Cell Transcriptome Atlas of the Aging Drosophila Brain. Cell 174, 982–998.e20. 10.1016/j.cell.2018.05.057

Davis, K.C., Raizen, D.M., 2017. A mechanism for sickness sleep: lessons from invertebrates. The Journal of Physiology 595, 5415–5424. 10.1113/JP273009

De, J., Wu, M., Lambatan, V., Hua, Y., Joiner, W.J., 2023. Re-examining the role of the dorsal fan-shaped body in promoting sleep in *Drosophila*. Current Biology 33, 3660–3668.e4. 10.1016/j.cub.2023.07.043

Dewa, K.-I., Kaseda, K., Kuwahara, A., Kubotera, H., Yamasaki, A., Awata, N., Komori, A., Holtz, M.A., Kasai, A., Skibbe, H., Takata, N., Yokoyama, T., Tsuda, M., Numata, G., Nakamura, S., Takimoto, E., Sakamoto, M., Ito, M., Masuda, T., Nagai, J., 2025. The astrocytic ensemble acts as a multiday trace to stabilize memory. Nature 648, 146–156. 10.1038/s41586-025-09619-2

Doherty, J., Logan, M.A., Taşdemir, O.E., Freeman, M.R., 2009. Ensheathing glia function as phagocytes in the adult Drosophila brain. J Neurosci 29, 4768–4781. 10.1523/JNEUROSCI.5951-08.2009

Dopp, J., Ortega, A., Davie, K., Poovathingal, S., Baz, E.-S., Liu, S., 2024. Single-cell transcriptomics reveals that glial cells integrate homeostatic and circadian processes to drive sleep-wake cycles. Nat Neurosci 27, 359–372. 10.1038/s41593-023-01549-4

Fernandes, V.M., Auld, V., Klämbt, C., 2024. Glia as Functional Barriers and Signaling Intermediaries. Cold Spring Harb Perspect Biol 16, a041423. 10.1101/cshperspect.a041423

Flores-Valle, A., Vishniakou, I., Seelig, J.D., 2025. Dynamics of glia and neurons regulate homeostatic rest, sleep and feeding behavior in Drosophila. Nat Neurosci 28, 1226–1240. 10.1038/s41593-025-01942-1

Freeman, M.R., Doherty, J., 2006. Glial cell biology in Drosophila and vertebrates. Trends Neurosci 29, 82–90. 10.1016/j.tins.2005.12.002

Guttenplan, K.A., Maxwell, I., Santos, E., Borchardt, L.A., Manzo, E., Abalde-Atristain, L., Kim, R.D., Freeman, M.R., 2025. GPCR signaling gates astrocyte responsiveness to neurotransmitters and control of neuronal activity. Science 388, 763–768. 10.1126/science.adq5729

Handy, G., Borisyuk, A., 2023. Investigating the ability of astrocytes to drive neural network synchrony. PLoS Comput Biol 19, e1011290. 10.1371/journal.pcbi.1011290

Hauglund, N.L., Andersen, M., Tokarska, K., Radovanovic, T., Kjaerby, C., Sørensen, F.L., Bojarowska, Z., Untiet, V., Ballestero, S.B., Kolmos, M.G., Weikop, P., Hirase, H., Nedergaard, M., 2025. Norepinephrine-mediated slow vasomotion drives glymphatic clearance during sleep. Cell 188, 606–622.e17. 10.1016/j.cell.2024.11.027

Haynes, P.R., Pyfrom, E.S., Li, Y., Stein, C., Cuddapah, V.A., Jacobs, J.A., Yue, Z., Sehgal, A., 2024. A neuron–glia lipid metabolic cycle couples daily sleep to mitochondrial homeostasis. Nat Neurosci 1–13. 10.1038/s41593-023-01568-1

He, T., Nitabach, M.N., Lnenicka, G.A., 2018. Parvalbumin expression affects synaptic development and physiology at the Drosophila larval NMJ. J Neurogenet 32, 209–220. 10.1080/01677063.2018.1498496

Huang, C., Maxey, J.R., Sinha, S., Savall, J., Gong, Y., Schnitzer, M.J., 2018. Long-term optical brain imaging in live adult fruit flies. Nat Commun 9, 872. 10.1038/s41467-018-02873-1

Ilanges, A., Shiao, R., Shaked, J., Luo, J.-D., Yu, X., Friedman, J.M., 2022. Brainstem ADCYAP1+ neurons control multiple aspects of sickness behaviour. Nature 609, 761–771. 10.1038/s41586-022-05161-7

Ingiosi, A.M., Hayworth, C.R., Harvey, D.O., Singletary, K.G., Rempe, M.J., Wisor, J.P., Frank, M.G., 2020. A Role for Astroglial Calcium in Mammalian Sleep and Sleep Regulation. Curr Biol 30, 4373–4383.e7. 10.1016/j.cub.2020.08.052

Irwin, M.R., 2019. Sleep and inflammation: partners in sickness and in health. Nat Rev Immunol 19, 702–715. 10.1038/s41577-019-0190-z

Jin, H., Li, M., Jeong, E., Castro-Martinez, F., Zuker, C.S., 2024. A body-brain circuit that regulates body inflammatory responses. Nature 630, 695–703. 10.1038/s41586-024-07469-y

Jones, J.D., Holder, B.L., Montgomery, A.C., McAdams, C.V., He, E., Burns, A.E., Eiken, K.R., Vogt, A., Velarde, A.I., Elder, A.J., McEllin, J.A., Dissel, S., 2025. The dorsal fan-shaped body is a neurochemically heterogeneous sleep-regulating center in Drosophila. PLOS Biology 23, e3003014. 10.1371/journal.pbio.3003014

Kent, S., Bluthé, R.M., Kelley, K.W., Dantzer, R., 1992. Sickness behavior as a new target for drug development. Trends Pharmacol Sci 13, 24–28. 10.1016/0165-6147(92)90012-u

Koh, K., Joiner, W.J., Wu, M.N., Yue, Z., Smith, C.J., Sehgal, A., 2008. Identification of SLEEPLESS, a sleep-promoting factor. Science 321, 372–376. 10.1126/science.1155942

Konsman, J.P., Parnet, P., Dantzer, R., 2002. Cytokine-induced sickness behaviour: mechanisms and implications. Trends in Neurosciences 25, 154–159. 10.1016/S0166-2236(00)02088-9

Kremer, M.C., Jung, C., Batelli, S., Rubin, G.M., Gaul, U., 2017. The glia of the adult Drosophila nervous system. Glia 65, 606–638. 10.1002/glia.23115

Kume, K., Kume, S., Park, S.K., Hirsh, J., Jackson, F.R., 2005. Dopamine is a regulator of arousal in the fruit fly. J Neurosci 25, 7377–7384. 10.1523/JNEUROSCI.2048-05.2005

Kuo, T.-H., Handa, A., Williams, J.A., 2012. Quantitative Measurement of the Immune Response and Sleep in Drosophila. J Vis Exp 4355. 10.3791/4355

Kuo, T.-H., Pike, D.H., Beizaeipour, Z., Williams, J.A., 2010. Sleep triggered by an immune response in Drosophila is regulated by the circadian clock and requires the NFkappaB Relish. BMC Neurosci 11, 17. 10.1186/1471-2202-11-17

Lee, S.-H., Dan, Y., 2012. Neuromodulation of brain states. Neuron 76, 209–222. 10.1016/j.neuron.2012.09.012

Lefton, K.B., Wu, Y., Dai, Y., Okuda, T., Zhang, Y., Yen, A., Rurak, G.M., Walsh, S., Manno, R., Myagmar, B.-E., Dougherty, J.D., Samineni, V.K., Simpson, P.C., Papouin, T., 2025. Norepinephrine signals through astrocytes to modulate synapses. Science 388, 776–783. 10.1126/science.adq5480

Lenz, O., Xiong, J., Nelson, M.D., Raizen, D.M., Williams, J.A., 2015. FMRFamide signaling promotes stress-induced sleep in Drosophila. Brain Behav Immun 47, 141–148. 10.1016/j.bbi.2014.12.028

Li, H., Janssens, J., De Waegeneer, M., Kolluru, S.S., Davie, K., Gardeux, V., Saelens, W., David, F.P.A., Brbić, M., Spanier, K., Leskovec, J., McLaughlin, C.N., Xie, Q., Jones, R.C., Brueckner, K., Shim, J., Tattikota, S.G., Schnorrer, F., Rust, K., Nystul, T.G., Carvalho-Santos, Z., Ribeiro, C., Pal, S., Mahadevaraju, S., Przytycka, T.M., Allen, A.M., Goodwin, S.F., Berry, C.W., Fuller, M.T., White-Cooper, H., Matunis, E.L., DiNardo, S., Galenza, A., O’Brien, L.E., Dow, J.A.T., FCA Consortium§, Jasper, H., Oliver, B., Perrimon, N., Deplancke, B., Quake, S.R., Luo, L., Aerts, S., Agarwal, D., Ahmed-Braimah, Y., Arbeitman, M., Ariss, M.M., Augsburger, J., Ayush, K., Baker, C.C., Banisch, T., Birker, K., Bodmer, R., Bolival, B., Brantley, S.E., Brill, J.A., Brown, N.C., Buehner, N.A., Cai, X.T., Cardoso-Figueiredo, R., Casares, F., Chang, A., Clandinin, T.R., Crasta, S., Desplan, C., Detweiler, A.M., Dhakan, D.B., Donà, E., Engert, S., Floc’hlay, S., George, N., González-Segarra, A.J., Groves, A.K., Gumbin, S., Guo, Y., Harris, D.E., Heifetz, Y., Holtz, S.L., Horns, F., Hudry, B., Hung, R.-J., Jan, Y.N., Jaszczak, J.S., Jefferis, G.S.X.E., Karkanias, J., Karr, T.L., Katheder, N.S., Kezos, J., Kim, A.A., Kim, S.K., Kockel, L., Konstantinides, N., Kornberg, T.B., Krause, H.M., Labott, A.T., Laturney, M., Lehmann, R., Leinwand, S., Li, J., Li, J.S.S., Li, Kai, Li, Ke, Li, L., Li, T., Litovchenko, M., Liu, H.-H., Liu, Y., Lu, T.-C., Manning, J., Mase, A., Matera-Vatnick, M., Matias, N.R., McDonough-Goldstein, C.E., McGeever, A., McLachlan, A.D., Moreno-Roman, P., Neff, N., Neville, M., Ngo, S., Nielsen, T., O’Brien, C.E., Osumi-Sutherland, D., Özel, M.N., Papatheodorou, I., Petkovic, M., Pilgrim, C., Pisco, A.O., Reisenman, C., Sanders, E.N., Dos Santos, G., Scott, K., Sherlekar, A., Shiu, P., Sims, D., Sit, R.V., Slaidina, M., Smith, H.E., Sterne, G., Su, Y.-H., Sutton, D., Tamayo, M., Tan, M., Tastekin, I., Treiber, C., Vacek, D., Vogler, G., Waddell, S., Wang, W., Wilson, R.I., Wolfner, M.F., Wong, Y.-C.E., Xie, A., Xu, J., Yamamoto, S., Yan, J., Yao, Z., Yoda, K., Zhu, R., Zinzen, R.P., 2022. Fly Cell Atlas: A single-nucleus transcriptomic atlas of the adult fruit fly. Science 375, eabk2432. 10.1126/science.abk2432

Liu, Y., Huang, J., Pandey, R., Liu, P., Therani, B., Qiu, Q., Rao, S., Geurts, A.M., Cowley, A.W., Greene, A.S., Liang, M., 2023. Robustness of single-cell RNA-seq for identifying differentially expressed genes. BMC Genomics 24, 371. 10.1186/s12864-023-09487-y

Ma, Z., Stork, T., Bergles, D.E., Freeman, M.R., 2016. Neuromodulators signal through astrocytes to alter neural circuit activity and behaviour. Nature 539, 428–432. 10.1038/nature20145

Marvel, F.A., Chen, C.-C., Badr, N., Gaykema, R.P.A., Goehler, L.E., 2004. Reversible inactivation of the dorsal vagal complex blocks lipopolysaccharide-induced social withdrawal and c-Fos expression in central autonomic nuclei. Brain Behav Immun 18, 123–134. 10.1016/j.bbi.2003.09.004

Measuring Sleep in Drosophila | Springer Nature Experiments [WWW Document], n.d. URL https://experiments.springernature.com/articles/10.1007/978-1-0716-2321-3_4 (accessed 1.28.26).

Miyashita, T., Murakami, K., Kikuchi, E., Ofusa, K., Mikami, K., Endo, K., Miyaji, T., Moriyama, S., Konno, K., Muratani, H., Moriyama, Y., Watanabe, M., Horiuchi, J., Saitoe, M., 2023. Glia transmit negative valence information during aversive learning in Drosophila. Science 382, eadf7429. 10.1126/science.adf7429

Nakagawa, H., Maehara, S., Kume, K., Ohta, H., Tomita, J., 2022. Biological functions of α2-adrenergic-like octopamine receptor in Drosophila melanogaster. Genes, Brain and Behavior 21, e12807. 10.1111/gbb.12807

Oliveira, J.F., Agarwal, A., Beckervordersandforth, R., Curreli, S., Denizot, A., Dulla, C., Enger, R., Goda, Y., Goshen, I., Holt, M.G., Nimmerjahn, A., Perea, G., Scimemi, A., Henneberger, C., 2026. The multiple scales of astrocytic functional units. Nat Neurosci 29, 1279–1292. 10.1038/s41593-026-02308-x

Orthmann-Murphy, J.L., Freidin, M., Fischer, E., Scherer, S.S., Abrams, C.K., 2007. Two distinct heterotypic channels mediate gap junction coupling between astrocyte and oligodendrocyte connexins. J Neurosci 27, 13949–13957. 10.1523/JNEUROSCI.3395-07.2007

Osorio-Forero, A., Foustoukos, G., Cardis, R., Cherrad, N., Devenoges, C., Fernandez, L.M.J., Lüthi, A., 2025. Infraslow noradrenergic locus coeruleus activity fluctuations are gatekeepers of the NREM-REM sleep cycle. Nat Neurosci 28, 84–96. 10.1038/s41593-024-01822-0

Osterhout, J.A., Kapoor, V., Eichhorn, S.W., Vaughn, E., Moore, J.D., Liu, D., Lee, D., DeNardo, L.A., Luo, L., Zhuang, X., Dulac, C., 2022a. A preoptic neuronal population controls fever and appetite during sickness. Nature 606, 937–944. 10.1038/s41586-022-04793-z

Osterhout, J.A., Kapoor, V., Eichhorn, S.W., Vaughn, E., Moore, J.D., Liu, D., Lee, D., DeNardo, L.A., Luo, L., Zhuang, X., Dulac, C., 2022b. A preoptic neuronal population controls fever and appetite during sickness. Nature 606, 937–944. 10.1038/s41586-022-04793-z

Otto, N., Marelja, Z., Schoofs, A., Kranenburg, H., Bittern, J., Yildirim, K., Berh, D., Bethke, M., Thomas, S., Rode, S., Risse, B., Jiang, X., Pankratz, M., Leimkühler, S., Klämbt, C., 2018. The sulfite oxidase Shopper controls neuronal activity by regulating glutamate homeostasis in Drosophila ensheathing glia. Nat Commun 9, 3514. 10.1038/s41467-018-05645-z

Peng, H.-R., Zhang, Y.-K., Zhou, J.-W., 2023. The Structure and Function of Glial Networks: Beyond the Neuronal Connections. Neurosci Bull 39, 531–540. 10.1007/s12264-022-00992-w

Poskanzer, K.E., Yuste, R., 2016. Astrocytes regulate cortical state switching in vivo. Proc Natl Acad Sci U S A 113, E2675–2684. 10.1073/pnas.1520759113

Pyfrom, E.S., Beveridge, C., Haynes, P.R., Kanigicherla, V.A., Randolph, C.E., Costa, P.C., Negatu, S.G., Iyer, S., Killiany, S.L., Yue, Z., Astacio, E.N., Walker, K.A., Luu, K.N., Pivarshev, P.A., Jurado, K.A., Chopra, G., Sehgal, A., 2025. Neutral lipid processing in glia is sexually dimorphic and promotes sleep through diacylglycerol catabolism. 10.1101/2025.09.10.674993

Rash, J.E., Yasumura, T., Dudek, F.E., Nagy, J.I., 2001. Cell-specific expression of connexins and evidence of restricted gap junctional coupling between glial cells and between neurons. J Neurosci 21, 1983–2000. 10.1523/JNEUROSCI.21-06-01983.2001

Sánchez Romero, J., Navarrete, M., 2026. Astroengrams: rethinking the cellular substrate for memory. Nat Rev Neurosci 27, 289–300. 10.1038/s41583-025-01012-2

Schneider, A., Ruppert, M., Hendrich, O., Giang, T., Ogueta, M., Hampel, S., Vollbach, M., Büschges, A., Scholz, H., 2012. Neuronal Basis of Innate Olfactory Attraction to Ethanol in Drosophila. PLOS ONE 7, e52007. 10.1371/journal.pone.0052007

Silverman, D., Chen, C., Chang, S., Bui, L., Zhang, Y., Raghavan, R., Jiang, A., Le, A., Darmohray, D., Sima, J., Ding, X., Li, B., Ma, C., Dan, Y., 2025. Activation of locus coeruleus noradrenergic neurons rapidly drives homeostatic sleep pressure. Sci Adv 11, eadq0651. 10.1126/sciadv.adq0651

Singh, P., Donlea, J.M., 2020. Bidirectional Regulation of Sleep and Synapse Pruning after Neural Injury. Curr Biol 30, 1063–1076.e3. 10.1016/j.cub.2019.12.065

Stahl, B.A., Peco, E., Davla, S., Murakami, K., Caicedo Moreno, N.A., van Meyel, D.J., Keene, A.C., 2018. The Taurine Transporter Eaat2 Functions in Ensheathing Glia to Modulate Sleep and Metabolic Rate. Curr Biol 28, 3700–3708.e4. 10.1016/j.cub.2018.10.039

Suri, V., Qian, Z., Hall, J.C., Rosbash, M., 1998. Evidence that the TIM light response is relevant to light-induced phase shifts in Drosophila melanogaster. Neuron 21, 225–234. 10.1016/s0896-6273(00)80529-2

Theparambil, S.M., Begum, G., Rose, C.R., 2024. pH regulating mechanisms of astrocytes: A critical component in physiology and disease of the brain. Cell Calcium 120, 102882. 10.1016/j.ceca.2024.102882

Troha, K., Im, J.H., Revah, J., Lazzaro, B.P., Buchon, N., 2018. Comparative transcriptomics reveals CrebA as a novel regulator of infection tolerance in D. melanogaster. PLoS Pathog 14, e1006847. 10.1371/journal.ppat.1006847

Vanderheyden, W.M., Goodman, A.G., Taylor, R.H., Frank, M.G., Van Dongen, H.P.A., Gerstner, J.R., 2018. Astrocyte expression of the Drosophila TNF-alpha homologue, Eiger, regulates sleep in flies. PLoS Genet 14, e1007724. 10.1371/journal.pgen.1007724

Watkins, L.R., Maier, S.F., Goehler, L.E., 1995. Cytokine-to-brain communication: a review & analysis of alternative mechanisms. Life Sci 57, 1011–1026. 10.1016/0024-3205(95)02047-m

Yildirim, K., Petri, J., Kottmeier, R., Klämbt, C., 2019. Drosophila glia: Few cell types and many conserved functions. Glia 67, 5–26. 10.1002/glia.23459

Zhang, J., Brown, E.B., Lloyd, E., Yeragi, E., Farhy-Tselnicker, I., Keene, A.C., 2026. Sleep rescues age-associated loss of glial engulfment. PLoS Genet 22, e1011999. 10.1371/journal.pgen.1011999

Zhang, Y., Noya, S.B., Li, Y., Fang, J., Sehgal, A., 2025. The microbiome interacts with the circadian clock and dietary composition to regulate metabolite cycling in the Drosophila gut. Elife 13, RP97130. 10.7554/eLife.97130

Zhu, R., Khorbtli, S., Zhang, J., Fu, Z., Huang, C., 2026. Protocol for a saline-free surgical preparation of adult Drosophila for chronic in vivo brain imaging. STAR Protocols 7. 10.1016/j.xpro.2026.104550

